# Breaking dormancy in spores of budding yeast transforms its cytoplasm and the solubility of its proteome

**DOI:** 10.1101/2022.07.29.502016

**Authors:** Samuel Plante, Kyung-Mee Moon, Pascale Lemieux, Leonard J. Foster, Christian R. Landry

**Affiliations:** Institut de Biologie Intégrative et des Systèmes (IBIS), 1030, avenue de la Médecine, Université Laval, Québec (Québec), Canada, G1V 0A6; Regroupement Québécois de Recherche sur la Fonction, l’Ingénierie et les Applications des Protéines, (PROTEO), 1045 Avenue de la Médecine, Université Laval, Québec (Québec), Canada, G1V 0A6; Département de biologie, 1045 Avenue de la Médecine, Université Laval, Québec (Québec), Canada, G1V 0A6.; Département de biochimie, microbiologie et bio-informatique, 1045 Avenue de la Médecine, Université Laval, Québec (Québec), Canada, G1V 0A6.; Centre de recherche en données massives (CRDM), 1065, avenue de la Médecine, Université Laval, Québec (Québec), Canada, G1V 0A6; Department of Biochemistry & Molecular Biology, and Michael Smith Laboratories, University of British Columbia, Vancouver, BC, Canada, V6T 1Z4

**Keywords:** cytoplasm, cell dormancy, protein solubility, protein phosphorylation

## Abstract

The biophysical properties of the cytoplasm are major determinants of key cellular processes and adaptation. Many yeasts produce dormant spores that can withstand extreme conditions. We show that spores of *Saccharomyces cerevisiæ* exhibit extraordinary biophysical properties, including a highly viscous and acidic cytosol. These conditions alter the solubility of more than 100 proteins such as metabolic enzymes that become more soluble as spores transit to active cell proliferation upon nutrient repletion. A key regulator of this transition is the heat shock protein Hsp42, which shows transient solubilization and phosphorylation, and is essential for the transformation of the cytoplasm during germination. Germinating spores therefore return to growth through the dissolution of protein assemblies, orchestrated in part by Hsp42 activity. The modulation of spores’ molecular properties are likely key adaptive features of their exceptional survival capacities.

## Introduction

Organisms across the tree of life rely on dormancy to withstand hostile conditions. This cellular state implies an arrest of the cell cycle and of cell metabolism, and changes in cell properties that favor survival under unfavorable conditions (Gremer and Sala 2013; Miller et al. 2021). For instance, nematodes, rotifers and tardigrades produce dormant life-stages that allow them to resist acute stresses such as freezing, desiccation and heat stresses (Guidetti, Altiero, and Rebecchi 2011; García-Roger et al. 2019; Vlaar et al. 2021). In flowering plants, the embryo develops as a dormant seed, which contributes to its survival over a long period of time by resisting drought and mechanical stress until it reaches favorable conditions to resume growth (Steven Penfield 2017). Cell dormancy is also an adaptive strategy in cancer cells, whereby metastatic cells become dormant after dissemination and resume proliferation after treatment has succeeded at eliminating the primary tumours (Phan and Croucher 2020). As one of the most widespread adaptive survival strategies to extreme conditions, understanding the molecular and cellular bases of cell dormancy is a major goal in cell biology.

Fungal life cycles include the production of spores. Although being formed through largely different mechanisms, conidia (asexual spores), and ascospores and basidiospores (sexual spores) have in common to be stable dormant cell types (Jan Dijksterhuis 2019; Wyatt, Wösten, and Dijksterhuis 2013). These cells all show a variety of resistance to extreme conditions such as heat, desiccation (Ho and Miller 1978; Oneto et al. 2020), and many harsh complex environment such as insect guts (Coluccio et al. 2008) or immune system assaults (Levitz and Diamond 1985; Botts and Hull 2010). Because of the increased resistance of spores to extreme conditions, sporulation is thought to be an adaptive strategy to survive changing environmental conditions (Huang and Hull 2017). In ascospores, which are produced by our model the budding yeast S. cerevisiae , stress resistance is largely attributed to the thick cell wall of specific composition (Neiman 2011), and to the accumulation of protective compounds like trehalose or mannitol (Kane and Roth 1974; Wyatt, Wösten, and Dijksterhuis 2013). These protective features develop during sporulation, which is typically induced in vegetative yeast by nutrient stress. When spores are exposed to favorable conditions, germination coordinates the breaking of dormancy and the loss of these protective features, with cell-cycle progression and vegetative growth resumption. This transition involves multiple changes in cellular state (Herman and Rine 1997), including the reactivation of multiple metabolic reactions. Although the precise nutrient stimuli that drive germination is dependent on ecological contexts, a carbon source such as glucose is typically an essential signal (Plante and Landry 2020b).

Recent studies have shown the potential complex influence of the physical properties and organization of the cytosol in dormancy and stress resistance. Cytosol’s viscosity, pH, crowding and protein phase separation have been linked to global cell adaptation across taxonomic groups. For instance, in tardigrades, desiccation resistance is mediated by intrinsically disordered proteins that form vitrified structures (Boothby et al. 2017). Seeds of the plant *Arabidopsis thaliana* sense hydration as the key trigger for their germination through phase separation of the protein Floe1 (Dorone et al. 2021). This process is a highly responsive environmental sensor since the biophysical state of Floe1 changes within minutes when water content is altered (Steve Penfield 2021). Examples of the responsiveness of the biophysics of the cell cytoplasm also come from yeast, such as *S. cerevisiæ*, responding to acute stresses. Early heat shock response in yeast includes cytoplasm acidification (Triandafillou et al. 2020), viscosity adaptation (Persson, Ambati, and Brandman 2020), and protein phase separation (Iserman et al. 2020; Riback et al. 2017; Wallace et al. 2015). Heat shock response induces the expression of many heat shock proteins composed mainly of molecular chaperones (Parsell and Lindquist 1993) which act as a dispersal system for the heat-induced phase-separated protein condensates that promotes the rapid recovery from stress (Yoo et al. 2022).

Given that budding yeast spores are inherently resistant to stresses that are known to modify many biophysical features of the cytoplasm, we hypothesize that the spore cytoplasm has biophysical properties similar to cells exposed to acute stress and that these will dynamically change during early spore germination. Here, we therefore examine the biophysical properties of dormant budding yeast ascospores and the changes that occur during dormancy breaking to unveil the molecular processes which support this critical life-history cell transition. Our results reveal that dormant spore cytosol is highly rigid and acidic, and that breaking of dormancy is supported by the neutralization and increased fluidity of the cytoplasm. We used mass spectrometry to examine and perform proteome wide measurements of protein solubility through germination. The measurements of 895 proteins revealed dynamic changes in protein solubility through germination. We uncovered, for instance, the solubilization of several metabolic enzymes during this transition. Our results demonstrate that spores have exceptional biophysical properties and that many of the changes taking place in spores mimic what occurs in yeast experiencing stress relief. One major similarity is the implication of a small heat shock protein, Hsp42 in spore germination, which is essential for normal spore activation and whose activity is regulated by its phosphorylation.

## Results and discussion

### Spores have a dense cytoplasm and display a different ultra architecture that changes during germination

Spore germination is the transition of dormant spores toward metabolically active and dividing vegetative yeast cells. Spores and vegetative yeast differ in terms of morphology and this morphology gradually changes through time. Spores are spherical and highly light refractile (Figure 1A), and darken and start growing quickly after the initiation of germination which can be induced by transferring cells to rich media. The hallmark of the completion of germination and the return to vegetative growth is bud emergence, which occurs at about 6 hours after induction of germination (Figure 1A). Spores’ transition from high to low refractility correlates with the decrease in optical density at 595 nm (*A*_595_) of the pure spore culture (Plante and Landry 2020a), with the minimal values reached about 3 hours after induction (Figure 1B). Then subsequent growth leads to an increase of optical density. One of the adaptive features of spores is their resistance to heat. This feature is lost during germination. The quantification of heat-shock resistance during germination highlights a drastic cellular transition as early as one hour after induction, at which point resistance to thermal stress decreases and reaches levels that compare to that of vegetative yeast (Figure 1C). Taken together, these measurements define the time-frame and the major time-points can be used to examine the underlying cellular and molecular changes.

**Figure 1.**
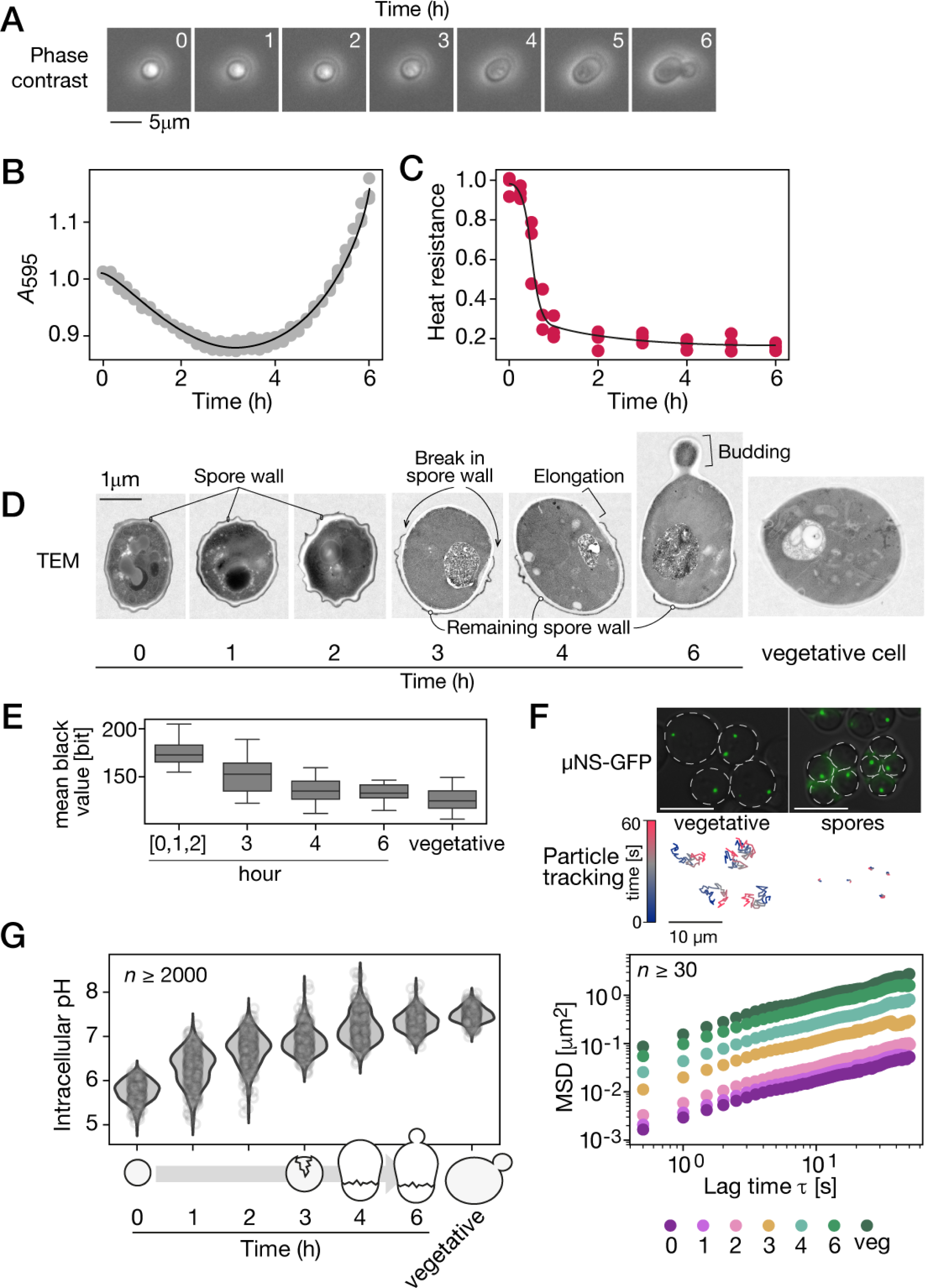
The cytoplasm of dormant spores displays high rigidity and density and is an acidic environment. A) Phase-contrast microscopic images of ascospore (same cell followed through time) at the indicated time after exposure to rich media, which activates germination. Scale bar represents 5 µm. B) Optical density (*A*_595_) and C) heat resistance of pure spore cultures through time after exposure to rich media. Heat resistance is the ratio of growth after a heat shock at 55°C for 10 minutes to growth without heat treatment. Experiments were performed in triplicate and values for individual replicates are shown. D) Representative transmission electron microscopic (TEM) images of spores at the indicated time after exposure to rich medium, and of a vegetatively growing yeast cell (vegetative). Cells were prepared and stained at the same time. Imaging was performed on a single layer. The scale bar represents 1 µm. See Figure S1A for more examples. E) Mean black level of spore cytosol at the indicated time after exposure to rich medium and of vegetative yeast. Spores at 0, 1 and 2-hour time-points are merged into a single category since they are indistinguishable from one another. F) Top, microscopic images of µNS-GFP particles in vegetative yeasts and spores. Underneath is the corresponding 1 minute trajectories of the particles. Color indicates time scale. Scale bar represents 10 µm. Bottom, ensemble Mean Squared Displacement (MSD) of µNS-GFP particles in spores at the indicated time after exposure to rich medium and in vegetative cells. G) Intracellular pH measured at the indicated time point after germination induction, and in exponentially growing cells. Measurements in at least 2000 cells are shown at each time point.

We obtained a more detailed view of the inner cell during germination using transmission electron microscopy (TEM, Figure 1D). Dormant spores are distinguishable by their small size and the thick spore wall, which is not seen in yeast. The spore cytoplasm appears darker in TEM in comparison to a vegetative yeast, which suggests a denser cytosol (Figure 1D, 1E, S1A). Spores have a different cytoplasmic organization. This is shown by the membranous structures that look highly packed in dormant spores compared to in vegetative yeast (Figure 1D). Cells at one and two hours into germination are still indistinguishable from dormant cells. Visible cytoplasmic organization changed after about three hours of germination, which correlated with a drop in heat resistance comparable to levels seen in vegetatively growing yeast (Figure 1D). At this time point, there is a rupture of the outer spore wall and the cell starts increasing in size where the spore wall is open (Figure 1D). This size increase is accompanied with a decrease in cytoplasm density (Figure 1E). These observations suggest that transformation of the physical nature of the cytosol environment coincides with germination and return to vegetative growth.

To test the cytosol physical properties during germination, we quantified its dynamic through the examination of macromolecular motion. We expressed the reovirus non-structural protein µNS tagged with GFP as a foreign tracer particle, which has shown to be a suited probe for subcellular environment in yeast (Munder et al. 2016).

µNS self-assembles in one or two discrete particles in the yeast cytoplasm that we could detect in both spores and vegetative yeast (Figure 1F, S2). The tracking of single particles revealed their lower mobility in dormant spores compared to vegetative yeast (Figure 1F). These measurements suggest that dormant spore cytoplasm is highly rigid or dense. Particle motion remained low during the first two hours of germination, then increased gradually from hatching (3 hour time-point) until the end of germination (Figure 1F). At bud emergence, the motion of µNS particles is close to that measured in vegetative yeast. Other experiments using µNS as tracer particles reported mean squared displacement (MSD) at 1 second lag-time in the order of 10^-1^ µm^2^ (Munder et al. 2016), which is in the range of our results. While energy deprivation in yeast reduces MSD of tracer particles by less than one order of magnitude (Munder et al. 2016; Joyner et al. 2016), we reported that particles have motion two orders of magnitude lower than in vegetative yeast. These results highlight that dormant spores have an exceptionally dense and rigid cytosol. These observations are in agreement with previous work on the fungi *Talaromyces macrosporus*, where spores were found to be characterized by high viscosity (J. Dijksterhuis et al. 2007).

Stress response in yeast includes cytoplasm acidification that culminates with its rigidification (Munder et al. 2016), including during heat shock (Triandafillou et al. 2020). We therefore hypothesized that the high viscosity of the spore cytoplasm and heat shock resistance would be accompanied by a low pH that would increase during germination. To test this hypothesis, we constitutively expressed the pH biosensor superfold-pHluorin (Miesenböck, De Angelis, and Rothman 1998), in both vegetative yeast and spores after calibrating pHluorin fluorescence *in vivo*. We estimated pH to be around 5.9 in dormant spores, confirming previous reports (Aon et al. 1997; Barton et al. 1980). Over the course of germination, the cytosol is gradually neutralized (Figure 1F). As soon as one hour after exposure to rich media, median intracellular pH rises to 6.2 and it slowly increases until the end of the process (pH_i_ = 7.3). At this point, intracellular pH gets close to that measured in vegetatively growing cells (pH_i_ = 7.4). Previous works showed that acidification and alkylation of yeast cytosol causes reduction and increase of motility of µNS particles respectively, and that this effect happens quickly, in the scale of a few minutes (Munder et al. 2016). However, our results show that during germination, the kinetics of change in particle mobility is delayed compared to change of intracellular pH. Although germination involves physicochemical changes related to that seen in vegetative yeast recovering from stress, they are modulated in a germination specific manner. Our results reveal the contribution of possibly many other factors to changes in viscosity.

Altogether, these experiments show that extreme physicochemical conditions prevail in dormant spores compared to vegetative yeast, namely a highly rigid and acidic cytoplasm. These conditions are modulated during the germination and return to vegetative growth. These intracellular properties that change during germination can play a critical role in cellular function and organization as they are some of the determinants of protein phase separation (Persson, Ambati, and Brandman 2020). Protein phase separation was shown to underlie heat shock response in yeast and many other forms of stress responses during cell dormancy (Franzmann and Alberti 2019). We therefore hypothesized that proteins could have a different solubility in spores and that the modification of physicochemical properties during germination affect their solubility in a time-dependent fashion.

### Protein solubility changes during germination

We adopted a physical separation technique similar to the one used in the context of heat shock to measure biochemical changes in protein solubility proteome-wide in budding yeast (Wallace et al. 2015). Protein sedimentation was driven by ultracentrifugation, and protein partitioning between the pellet and supernatant fractions was quantified by liquid-chromatography-coupled tandem mass spectrometry (LC-MS/MS, Figure 2A). We measured the proportion of each protein that partitioned in the pellet fraction using P_index_ as a proxy for desolubilization in three biological replicates at 4 time-points during germination, and in vegetative yeasts. In total, we detected 24,559 unique peptides corresponding to 2,614 proteins across the experiments. We restricted our analysis to the 895 proteins with at least two unique peptides that were detected at every time-point to measure P_index_ (Table S3). Values for these 895 proteins range from 0 to 1. Zero indicates that the protein was detected only in the supernatant, and 1 indicates that the protein was detected only in the pellet. Proteins with low Pindex are referred to as soluble proteins, while proteins with high P_index_ as less soluble ones. Replicated measurements were strongly correlated (Figure S2A). Moreover, the solubility profile of endogenous proteins in the fractionned cell extracts revealed by western blot is similar to their Pindex trajectories (Figure S3).

**Figure 2.**
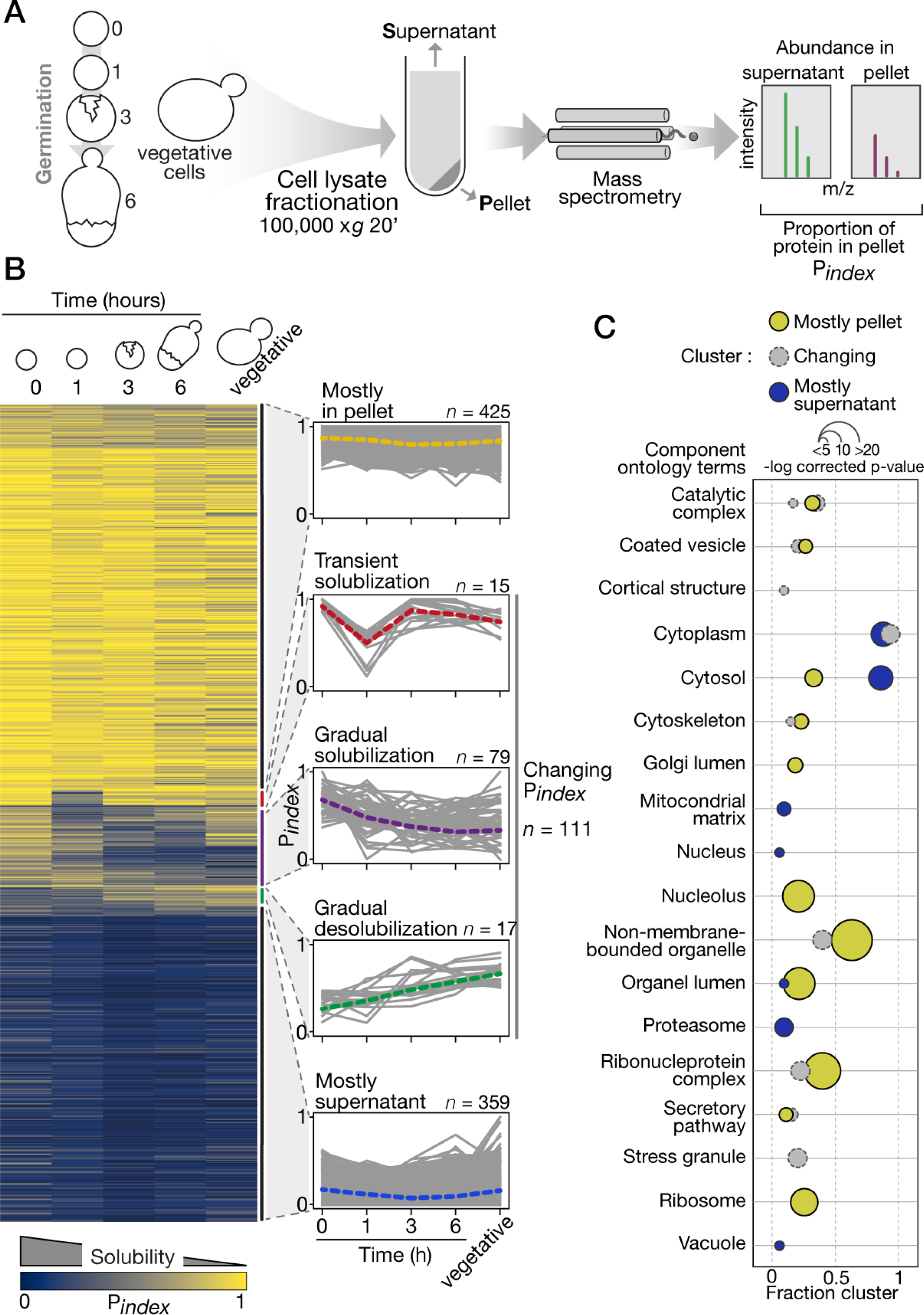
Proteome wide change in protein solubility during germination. A) Solubility measurement by LC-MS/MS estimates the proportion of each protein in the pellet (P_index_) at each major time-point sampled during germination. The experiment was performed in triplicate for all timepoints. B) Right, P_index_ values in the course of germination show, from top to bottom, proteins consistently found in the pellet, that transiently solubilize, that gradually solubilize, that gradually accumulate in the pellet, and that are consistently found in the supernatant. Left, individual P_index_ trajectories for each cluster determined by hierarchical clustering. The dotted line is the median trajectory for each cluster. C) Gene ontology term analysis focused on cellular component terms. Terms enriched for each cluster (Mostly in pellet, Changing Pindex and mostly supernatant) are shown as bubble plots. Colors refer to the cluster, position on the x-axis indicates the portion of the proteins in a cluster assigned to a GO term, and size of the bubble is scaled to the -log (*p*-values). See Figure S2 for additional details.

Five typical P_index_ trajectories were identified using hierarchical clustering (Figure 2B). The two largest clusters contain proteins that remain mostly soluble (mostly supernatant, n=359) and mostly insoluble (mostly in pellet, n=425). Together, they account for 87% of total proteins we considered in our analysis. This means that most of the proteins do not exhibit detectable changes in physicochemical partition during germination using our approach. However, 111 proteins showed changing P_index_ trajectories divided in three clusters. First, 15 proteins showed a transient solubilization early in germination. These proteins predominantly partitioned in the pellet in dormant spores, while one hour after exposure to rich media, their P_index_ dropped drastically before rising again at the three-hour time-point and remained insoluble until the end of germination. Another group of 17 proteins gradually desolubilize in the course of germination. They start with high solubility (low P_index_) in dormant spores, and gradually reach higher P_index_ value at later time-point in the process. Finally, 79 proteins with varying single trajectories gradually gained solubility during germination. Gene ontology analysis, using the total protein considered for our analysis as a reference set, revealed that clusters are enriched for different cell component terms (Figure 2C). While the mostly supernatant cluster appears to contain essentially cytosolic proteins, the proteins in the mostly pellet and changing Pindex clusters are assigned to various and more specific cellular components. For instance, the later clusters are both enriched for proteins in non-membrane-bounded-organelles, ribonucleoprotein complexes and cytoskeleton. There is a specific enrichment for nucleolar and ribosomal protein in the mostly pellet cluster, and a specific enrichment for stress granule proteins in the changing Pindex cluster. These results highlight the level of separation performed by our technique, which seems to separate non-membrane-bounded organelles and macromolecular complexes from the other constituents of the cytosol.

We examined the properties of proteins that associate with these changes in solubility. Proteins that change solubility are not more nor less abundant than other proteins (Figure S2B). Principal component analysis (PCA) revealed that of all the protein properties considered, propensity for condensate formation (PSAP, (van Mierlo et al. 2021)) and score for prion-like domains prediction (PLAAC, (Lancaster et al. 2014) ) are the ones that contribute the most to the separation of proteins in terms of P_index_ (Figure S2C). Because prion-like domains can contribute to protein phase-separation and tune the dynamics of biomolecular condensate (Holehouse et al. 2021), these results corroborate the enrichments for non-membrane-bounded organelles and macromolecular complexes we reported in the clusters mostly in pellet and changing P_index_. However, insolubility does not necessarily reflect phase separation as protein solubility is also influenced by many other factors such as misfolding, formation of protein/RNA granules, or other homogeneous or heterogeneous oligomerization. Nevertheless, because propensity for condensate formation positively correlates with P_index_, high P_index_ estimates at least partially reflect phase-separation of proteins and macromolecular assemblies. In addition, the analysis of known physical interaction among the detected proteins revealed that changes we reported in P*index* do correlate with large interaction networks. Out of the 111 changing Pindex proteins, 6 pairs of physically interacting proteins were found (Figure S4). This suggests that the changes in protein organization we observed likely reflect bulk changes in cytoplasm properties rather than remodeling of specific interactions.

### Many classes of proteins change solubility during germination, including metabolic enzymes

To understand the functional significance of change in P_index_, we searched for gene ontology (GO) terms enrichment in three clusters that display dynamic change. First, in the transient solubilization cluster we found significant enrichment for lipid and phospholipid binding proteins (Figure 3A). This group includes for instance the translation initiation factor Cdc33 and the GTP-binding protein Ras2 (Figure S5C). In the gradual desolubilization cluster, which includes for instance the transcription elongation factor Spt5 and the vacuolar carboxypeptidase Cps1, we found enrichment for phosphatidylinositol-3-phosphate (PI3P)-binding proteins (Figure 3A). We suspect that the modulation in solubility we detect in these clusters is a reflection of the gain of activity of many cellular pathways. For instance, gradual insolubility of the SNARE chaperone Sec18 may reflect increasing assembly of membrane-fusion complexes as vesicle transport is resumed to sustain cell growth. Finally, the gradual solubilization cluster is enriched for proteins involved in metabolic process: precisely, amino acid and carbohydrate metabolism, and protein phosphatase activity (Figure 3A), including for instance the ceramide-activated protein phosphatase Sit4 which functions in the G1/S transition in cell-cycle (Barbosa et al. 2016). This may reflect the reentry of the dormant spores in the cell cycle. Among this group, we also identified the stress related proteins Ola1 and Yef3, which are known to aggregate in response to heat stress and disaggregate during recovery (Wallace et al. 2015). The behaviour of these proteins suggest that dormancy in spores shares features with stress response and that germination would correspond to stress relief.

**Figure 3.**
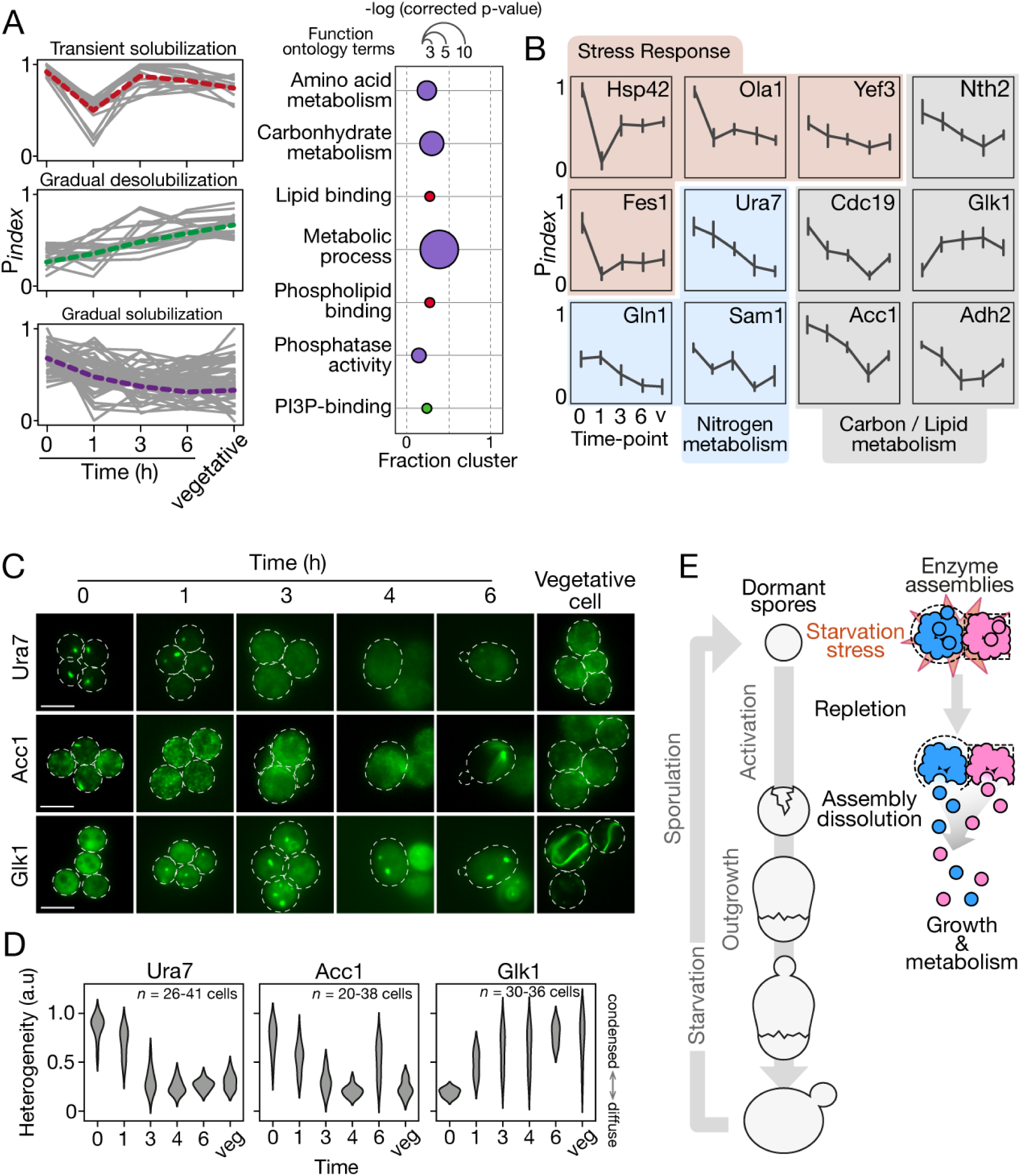
Solubility changes reflect metabolism activation and mimic stress relief during germination. A) Enrichment for gene ontology terms in each dynamically changing solubility cluster. Red, transiently solubilizing cluster; Green, gradual desolubilization cluster; Purple, gradual solubilization cluster. The position on the x-axis indicates the portion of the proteins in a cluster assigned to a GO term, and size of the bubble is scaled to the -log (*p*-values) from a hypergeometric test. B) Individual P_index_ trajectory for representative proteins through germination. Bar represents standard variation. Proteins are clustered by function; Red, stress response proteins; Blue, Nitrogen metabolism proteins; Gray, Lipid and carbon metabolism proteins. Error bars represent standard deviation of three replicates. C) Representative fluorescence microscopic images of spores expressing the indicated proteins tagged with GFP during germination. Top to bottom, Acetyl-CoA carboxylase Acc1 (Lipid biosynthesis), CTP synthase Ura7 (pyrimidines synthesis) and Glucokinase Glk1 (glycolysis). The Glk1 foci formation and the dissolution of Acc1 and Ura7 foci in course of germination support that dormancy in spores is analogous to a stress state and germination alleviates this state. Dotted lines indicate cell contour determined by brightfield images. Scale bars represent 5µm. D) Measure of cellular heterogeneity (coefficient of variation) of the fluorescent proteins in spore at the indicated time after exposure to rich medium or in vegetative cells. Between 20 and 41 cells were analyzed at each time-points. E) Schematics highlighting effects on protein solubility of nutrient starvation and repletion during sporulation and germination, respectively. Pink and blue assemblies represent assemblies of enzymes needed for growth and metabolism during dormancy, which disassemble (pink and blue circles) during germination.

Within the group of proteins with increasing solubility, we identified enzymes involved in carbohydrate, lipid and nitrogen metabolisms (Figure 3B). Since nutrient starvation is the key signal that triggers sporulation, the comportment of these metabolic enzymes that solubilize in the course of germination caught our interest. We investigated two of them: the CTP synthase Ura7 and the acetyl-CoA carboxylase Acc1, which are enzymes known to form high molecular weight assemblies in response to nutrient starvation (Petrovska et al. 2014; Narayanaswamy et al. 2009). To validate the solubility changes revealed by P_index_ trajectories, we generated cells expressing either Ura7 or Acc1 fused - at their genomic locus - with GFP. Both Ura7 and Acc1 formed cytoplasmic foci in dormant spores (Figure 3C, 3D). Upon germination, Ura7-GFP and Acc1-GFP fluorescence signals changed until they became mostly diffuse in dividing cells. This behavior confirms the dissolution of the protein assemblies observed in the P_index_ trajectories. In addition, we noted the opposite behavior of the glucokinase Glk1. Glk1’s P_index_ trajectory suggests it gains insolubility during germination (Figure 3C, 3D). Correspondingly, we found Glk1-GFP to be diffuse in dormant spores, then appears as dense assemblies in cells as soon as one hour after exposure to rich media and until the end of germination. Glk1 was found to polymerize and form filament during the transition from low to high sugar conditions (Stoddard et al. 2020). Its behavior in germinating cells again suggest that spores remain dormant in a starved form and that breaking of dormancy implies changes of enzyme biophysics in response to nutrient repletion.

The reverse order of events between spore germination and heat stress and nutrient stress responses for some key proteins suggests a model in which dormancy in spores is analogous to a stress response state and germination corresponds to the relief of the stress state allowing return to metabolic activity and vegetative growth (Figure 3D). Spores therefore most likely borrow stress resistance strategies we observe in vegetative yeast.

### The heat shock protein Hsp42 shows dynamic solubilization and phosphorylation during germination

To further explore the regulatory mechanisms driving cellular reorganization during germination, we searched in our proteomic data for phosphorylation on tyrosines, serines or threonines. We identified 36 phosphoproteins with a unique phosphopeptide in at least one time point during germination (Figure 4A). Given that we did not perform any enrichment for phosphorylation prior to mass spectrometry, the detection of a limited number of phosphorylation was expected. These include for instance the topoisomerase Top1 and the transcription elongation factor Spt5. One protein was at the intersection of the phosphoproteins cluster and the changing Pindex cluster (Figure 4B), namely the small oligomeric heat shock protein (sHSP) Hsp42. The small intersection between these clusters presuppose that phosphorylation is not a primary factor driving the change in protein solubility. Since stress response in vegetative yeast involves sHSP and they were recently identified as key players in the resolution of molecular assemblies that accompany heat shock (Yoo et al. 2022), we focused on this protein as a potential regulator of protein solubilization in germination.

**Figure 4.**
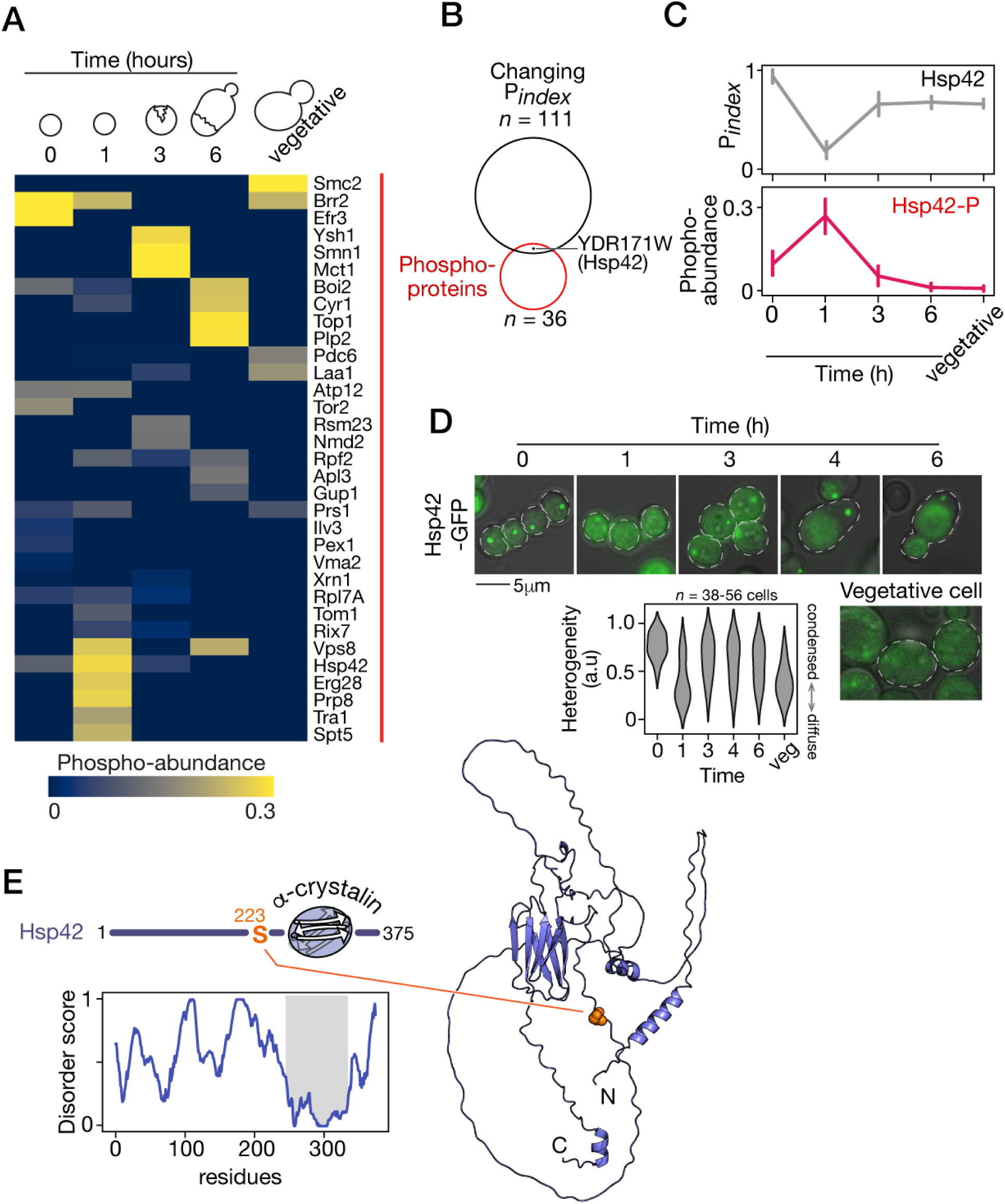
Hsp42 phosphorylation at S223 is synchronized with its transient solubilization. A) Relative abundance of the 36 phosphoproteins to the total abundance of each protein through germination. B) Hsp42 is phosphorylated during germination and changes solubility. See Figure S3 for additional information. C) Hsp42 is the only protein with dynamic solubility profile during germination that correlates with its dynamic phosphorylation, here at S223. Error bars represent standard deviation of three replicates. D) Representative fluorescence microscopic images of spores expressing Hsp42-GFP at the indicated time after the induction of germination. Dotted lines represent cell contour. Scale bar represents 5µm. Plot shows the cellular Hsp42-GFP heterogeneity score in spore at the indicated time after exposure to rich medium or in vegetative cells. E) Left, Disorder profile of Hsp42, predicted by Metapredict, shows the predicted structured ACD domain, and flanking disordered N- and C-terminal region. Right, predictedHsp42 structure. The S223 highlighted in orange is located in a disordered region.

Hsp42 is part of the protein clusters with changing solubility during germination. Furthermore, the solubility of Hsp42 is correlated with its phosphorylation during this time-period. Solubility transiently increases while abundance of its phosphorylation transiently increases (Figure 4B). Hsp42 was shown to reversibly assemble in heterogeneous granules in a heat-induced manner, or in quiescent cells in stationary phase (I.-C. Liu et al. 2012), and to function in tuning granules assembly and disassembly (Grousl et al. 2018). Remarkably, Hsp42-dependent spatial protein organization is crucial for cellular fitness, and lack in foci formation results in a significant delay when recovering from stationary phase (I.-C. Liu et al. 2012). Notably, sedimentation of Hsp42 was found to increase overtime in yeast faced with heat stress suggesting the trigger of its chaperon function, through a physical separation technique similar to ours (Wallace et al. 2015). We hypothesized that the dynamic in Hsp42 sedimentation we reported reflects its function during germination. Interestingly, Hsp42 has a unique behaviour among the molecular chaperone detected in our experiments (Figure S5D), which hints for an exclusive function.

To confirm the dynamic assembly and disassembly of Hsp42 during germination, we generated cells expressing Hsp42 fused to GFP. Hsp42 accumulates in cytoplasmic foci in dormant spores, which corroborates its solubility in the proteomics experiments. One hour after the induction of germination, Hsp42 is diffused, which shows the dissolution of the foci (Figure 4D). Diffused localization of Hsp42 is only transient since foci were visible at later time-point during vegetative growth. Microscopic observations therefore validate the P_index_ profile of Hsp42, suggesting a transient modification of the protein taking place early in germination.

The search for phosphorylation sites from the proteomics data revealed a dynamic phosphorylation site located in the N-terminal region (NTR) of Hsp42 (S223). Disorder profile of Hsp42 highlights three structurally distinct domains; a central structured domain that is predicted to a alpha-crystallin domain (ACD) common to sHSP, and a long N-terminal region (NTR) and a short C-terminal regions that are both predicted to be highly disordered (Martin Haslbeck, Weinkauf, and Buchner 2019). Structure prediction of Hsp42 (Uniprot Q12329) (Jumper et al. 2021) corroborates this architecture; it predicts with high confidence a beta-strand sandwich typical of ACD but predicts large unstructured parts in the N and C-terminal regions (Figure 4E). NTRs are shown to be involved in the regulation and dynamics of chaperone activity of sHSP (M. Haslbeck et al. 1999; Martin Haslbeck, Weinkauf, and Buchner 2019). These proteins are stored in an inactive form as high-order oligomers, and their activation involves phosphorylation, especially in the NTR, that drives disassembly of sHSP into smaller complexes. For instance, several phosphorylation sites on Hsp26 were found to activate chaperon activity by weakening interactions within the oligomers (Mühlhofer et al. 2021). The phosphorylation of S223 on Hsp42 has been previously detected by mass spectrometry. The abundance of this phosphorylation was found to increase in cells following exposure to heat (Kanshin et al. 2015). Hence, we hypothesized that the phosphorylation on S223 of Hsp42 is involved in this sHSP’s role during the major cytoplasmic changes that take place during germination.

### Hsp42 activity is crucial for normal progression of germination

We first confirmed that Hsp42 plays an important role in thermal stress protection, and tested if S223 phosphorylation may be regulating this function in vegetative yeast (Martin Haslbeck et al. 2004; Grousl et al. 2018). After being subjected to a heat shock, cells lacking Hsp42 (*hsp42*Δ::kanMX4) fail to grow as compared to WT cells, confirming thermal sensitivity (Figure 5A). A phosphomimetic mutant of Hsp42 (S223E) appears to be equally active as the WT chaperon, because expression of either protein tagged with GFP totally restores cellular heat shock resistance (Figure S6B). On the other hand, mutation of the site to a non-phosphorylatable residue (S223A) seems to impede chaperon activation or activity as revealed from the mutant phenotype. Cells expressing the Hsp42 S223A mutant show thermal stress sensitivity (Figure 5A) and this mutant fails to form large cytoplasmic foci as does the protective Hsp42 (Figure S6A). These results revealed that phosphorylation on this site is crucial for activation of Hp42 during heat stress. We therefore tested if Hsp42 activity was crucial during germination. Optical density decrease of *hsp42*Δ spores culture exposed to germination conditions is delayed compared to WT spores, suggesting a delay in germination (Figure 5C). Interestingly, microrheology revealed that *hsp42*Δ spores cytoplasm fluidify in a delayed fashion compared to WT spores (Figure 5D).

**Figure 5.**
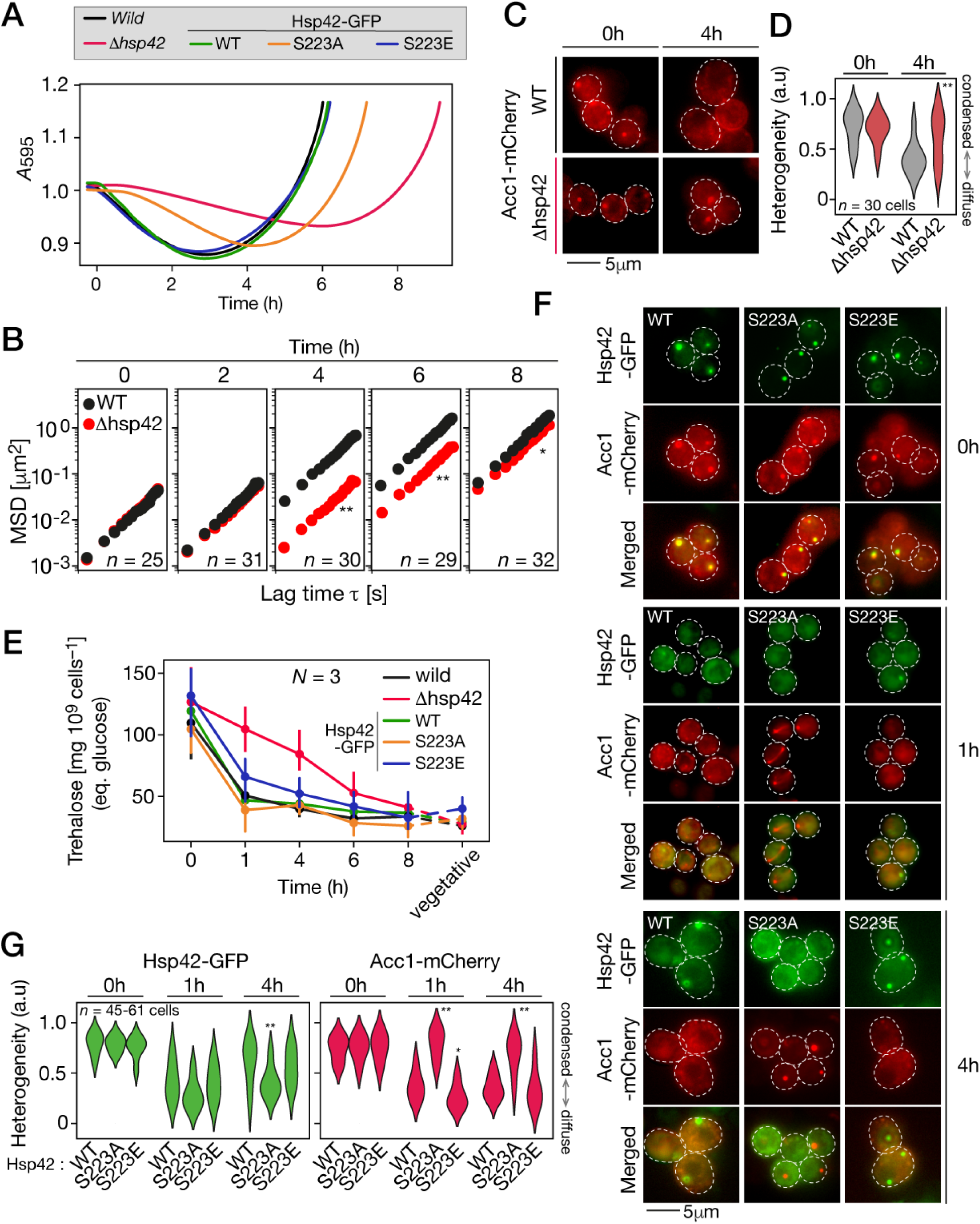
Active and phosphorylated Hsp42 is required for normal germination dynamics. A) Optical density of pure spore cultures of the indicated strains as a function of time following exposure to germination conditions. Shown are the mean values of three replicates. OD drop of the Δhsp42 spores is strongly delayed, indicating that germination is inhibited or slowed down. Spores expressing S223A Hsp42 mutant show a slight delay in germination, while germination of spores expressing S223E Hsp42 mutant is indistinguishable from that of wild spores. B) Ensemble mean square displacement of µNS-GFP in WT and Δhsp42 spores at the indicated time after exposure to rich medium. At each time point for each strain, 25 to 35 particles, corresponding to the same number of cells, were tracked. Kruskal-Wallis test, ** indicates p-value < 0.0001, * indicates p-value < 0.01. C) Fluorescence microscopy images of wild type (top) or ΔHsp42 spores expressing Acc1-mCherry at the indicated time after exposure to germination conditions. Scale bar represents 5µm. D) Cellular Acc1-mCherry heterogeneity score in WT or Δhsp42 spores at the indicated time after exposure to rich medium. E) Trehalose content (measured in equivalent of glucose concentration) in spores at the indicated time after exposure to rich medium, and in vegetative yeasts. Error bars represent standard deviation of three replicates. F) Fluorescence microscopy images of spores expressing either WT, S223A or S223E mutant Hsp42-GFP and Acc1-mCherry at the indicated time after exposure to germination conditions. Dotted lines represent cell contour. Scale bar represents 5 µm. Expression of S223A Hsp42-GFP mutant shows delay in Acc1 foci dissolution in spores.

Yeast adaptation to various stressor includes accumulation of compatible solute, notably the disaccharide trehalose (D’Amore, Crumplen, and Stewart 1991). It promotes survival by stabilizing macromolecules such as membranes and proteins (Kaushik and Bhat 2003; Magalhães et al. 2018). Fungal spores, including *S. cerevisiæ* ascospores, accumulate trehalose which account as a protective feature they develop (Kane and Roth 1974). Accumulation of trehalose was found to increase yeast cytoplasm viscosity as a homeostatic mechanism to maintain molecule diffusion rate in response to stress and energy depletion (Persson, Ambati, and Brandman 2020). Mobilization of trehalose is essential for ascospore germination since inhibition of trehalase activity in spores hinder germination (Thevelein, den Hollander, and Shulman 1984; Beltran et al. 2000). Here, we investigated trehalose content in the course of germination to see if it could play a role in the changes we measured. Both wild type and *hsp42*Δ spores accumulated a high level of trehalose. In wild type spores, trehalose content dropped by >55% after one hour in germination medium (Figure 5E), which is in line with previous works reporting a quick mobilization of trehalose content in the first hour of germination (Rousseau et al. 1972; Thevelein, den Hollander, and Shulman 1982). Contrastingly, trehalose content in hsp42Δ spores remained higher than the level in WT spores for the first four hours of germination, and then decreased down to WT level after 8 hours (Figure 5E). The delay in trehalose mobilization in *hsp42*Δ spore suggest that Hsp42 contributes to germination upstream of trehalose mobilization and that the major changes observed in the cytoplasm of *hsp42*Δ cells are at least partly caused by the dynamics of trehalose mobilization.

We tested how this delay in biophysical remodeling in *hsp42*Δ spores affected protein organization. In WT spores, Acc1-mCherry is diffusely localized four hours after exposure to rich media. In contrast, in *hsp42*Δ spores, Acc1-mCherry remains condensed as foci (Figure 5C, 5D). These results show a direct or indirect role for Hsp42 in disassembly of Acc1 foci. Altogether, our results reveal that presence of the chaperon Hsp42 is crucial for the remodeling of intracellular conditions taking place during germination.

Expression of either WT or a phosphomimetic mutant (S223E) of Hsp42 totally rescues the germination progression in *hsp42*Δ spores (Figure 5A) and restores the disassembly of Acc1 foci (Figure 5F, 5G). On the other hand, spores expressing the non-phosphorylatable S223A mutant experienced a delayed germination, and in these cells, Acc1 foci failed to disassemble (Figure 5F, 5G). Altogether, activation of Hsp42 through phosphorylation on S223 appears to be crucial to disassembly of low solubility protein and germination progression. Moreover, our results revealed a surprising contribution of phosphorylation to behavior of Hsp42 in germination. Both non-phosphorylatable S223A or phosphomimetic S223E Hsp42 mutants were solubilized at one-hour time-point, then at a later time-point (four hours), S223A mutant remained diffuse in the cytoplasm while WT and S223E mutant formed foci (Figure 5F). These results refine our hypothesis about the role of this phosphorylation on Hsp42 solubilization: they suggest that some processes independent of Hsp42 phosphorylation on S223 are involved. Among these processes is the quick mobilization of trehalose. Our results revealed the presence of Hsp42 is crucial for trehalose mobilization, but contrastingly, it occurs regardless of the Hsp42 mutant tested (Figure 5E). Altogether, our results highlight many functions of the molecular chaperone Hsp42 for germination progression, some of which involve its phosphorylation on S223.

## Conclusion

Some dormant cells have exceptional resistance to stress. What are the biophysical conditions that underlie these properties and how cells resume growth after dormancy are important questions across fields of biology. In this work, we used budding yeast ascospores to examine the biophysical properties of a dormant cytosol and its transition between dormancy and its return to vegetative growth. The spore cytosol is acidic and highly viscous, as has been observed in the context of various stresses, for instance in yeast cells during energy depletion (Munder et al. 2016), bacteria during metabolic arrest (Parry et al. 2014), dry plant seed (Buitink and Leprince 2008), and tardigrades during desiccation (Boothby et al. 2017). The properties observed in spores may therefore represent a conserved adaptive strategy for many cell types and species.

Because of the commonalities with the properties of yeast cells responding to stress, spores’ cytosolic properties reflect that spores are in metabolic repression and stress response state. During germination, cells come back to an unstressed state where spore cytosol is neutralized and its viscosity decreased. We also found massive altered protein organization in dormant spores that changes along with the cytosol pH and viscosity during germination. Germination therefore shares many features with stress relief, for instance after heat shock in yeast. One important question these observations trigger is what are the early molecular events that allow the cytosol and the solubility of many proteins to progressively change during germination. We identified Hsp42 as a key actor for the modulation of spore cytosol organization. The role of Hsp42 in dissolution of enzyme assembly during germination extends the role of chaperones in the disassembly of heat-induced protein condensate recently shown (Yoo et al. 2022). The role of heat shock proteins in response to stress in yeast may thus also be critical to the breaking of dormancy of spores, which have intrinsically high stress resistance. The dissolution of insoluble metabolic enzymes during this transition likely reflects the activation of spore metabolism as it modifies its physiology to respond to nutrients, and is an adaptation to nutrient repletion (Petrovska et al. 2014). We identified the phosphorylation of Hsp42 at S223 to be critical for its regulating function in protein organization. This post-translational modification has been reported multiple times in phosphoproteomic analysis (Swaney et al. 2013; Martínez-Montañés et al. 2020), notably in the context of heat shock where cells in unstressed conditions show low levels of Hsp42 phosphorylation while under stress conditions they quickly accumulate phosphorylated Hsp42 (Kanshin et al. 2015). Regarding these observations, the Hsp42 profile we report shows that early in germination spores exhibit stress response, and that while gemination progresses, stress response is relieved. The dynamic modulation in phosphorylation of Hsp42 we report implies that prior signaling steps including kinase activity upstream are required for the adaptation of spore cytosol organization to nutrient repletion. Therefore, Hsp42 presumably functions in adapting spore cytosol to nutrient repletion in a similar manner as Floe1 integrates the signal of adequate hydration in *A. thaliana* seed to control germination (Dorone et al. 2021). The first event involving Hsp42 is its solubilization occuring at the breaking of dormancy. The timing of Hsp42 foci dissolution at the onset of breaking of dormancy indicated that it is likely triggered by the pathways initiating germination. While this hypothesis is not substantiated in this work, it will be worth considering the mobilization of trehalose, which functions as a macromolecule stabilizer in dormant spores, as a factor that could interact with Hsp42 to modulate the solubility of the cytoplasm. Altogether, our results expand our knowledge of the molecular factor taking part in dissolution of protein assemblies, and sheds light into the regulation of protein condensate through signaling. Signaling and kinase activity has been previously linked with protein organization in the context of cellular stress (Wippich et al. 2013). Stress-induced phosphorylation of human Hsp27 was shown to cause its phase separation with FUS, a process that was found to prevent FUS amyloid fibril formation (Z. Liu et al. 2020). Phosphorylation of Hsp42 could imply the mitogen-activated protein (MAP) kinase signaling pathway, which has been reported to be involved in human Hsp27 phosphorylation (Z. Liu et al. 2020). Some kinases in yeast, notably cyclin-dependent kinases Cdc28 and Pho85 or MAP kinases Hog1 and Fus3, have specificities that correspond to the phosphorylation site motif of Hsp42 S223 (Mok et al. 2010). In addition, a target of the MAP kinase Hog1, the transcription elongation factor Spt5 (Silva et al. 2017), does show a similar profile of phosphorylation during germination (Figure 4A). Connecting upstream kinases to the activity of Hsp42 will eventually allow to connect nutrient sensing of activating spores and the biophysics of spore cytoplasm.

Cell dormancy is widespread across the tree of life and is a survival strategy for many species facing harsh conditions and pathogens (Ortiz, Huang, and Hull 2019) and cancer cells facing drug treatment (Oren 2022). By discovering what are the early events that regulate the breaking of dormancy, our work will help better understand the molecular basis of adaptation to extreme conditions and potentially help find ways to develop drugs or conditions that can potentiate existing drugs to overcome their exceptional resistance mechanisms.

## Acknowledgments

We thank Daniel Evans-Yamamoto, David Bradley and Alexandre K. Dubé for their comments on the manuscript, and Alexandre K. Dubé and Isabelle Gagnon-Arsenault for their support in the laboratory. We are grateful to Pr Martin Bisaillon, from Université de Sherbrooke (Canada) for providing us the µNS coding sequence. This work was supported by NSERC Discovery Grants to CRL and LJF, a Canadian Institutes of Health Research (CIHR) Foundation grant (387697) to CRL, and platform funding from Genome Canada (264PRO) to LJF. CRL holds the Canada Research Chair in Cellular Synthetic and Systems Biology.

## Authors contributions

Conceptualization: CRL and SP

Methodology and Experiments: SP, KMM, PL, LJF

Validation: SP

Formal analysis: SP, KMM

Writing – original draft preparation: SP, CRL

Writing – review and editing: all authors

Visualization: SP

Supervision: CRL and LJF

Project administration: CRL

Funding acquisition: CRL and LJF

## Methods

### Yeast strains construction and culture conditions

The yeast strains used in this study are listed in table S1. Background for every construction is the wild diploid *Saccharomyces cerevisiae* LL13_054 (Leducq et al. 2016). This strain was chosen for its propency to sporulate at high efficiency. For C-terminal labeling of Acc1, Ura7, Glk1 and Hsp42 with GFP at their native genomic locus (Figure 3C, 4D), GFP and Hyg resistance marker (hphNT2) were amplified from pYM25 with flanking DNA for genomic integration. For deletion of *HSP42* (Figure 5, *hsp42*Δ) the cassette loxP-pAgTEF1-kanMX-tAgTEF1-loxP from pUG6 was amplified with the flanking DNA for replacement of the HSP42 coding sequence, leaving its promoter and terminator intact. The deletion cassette was removed by expressing the recombinase Cre on the plasmid pNatCRE. Site-directed mutagenesis on Hsp42 (S223A or S223E) was conducted by primer extension. For restoration of Hsp42 expression in hsp42Δ cells (Figure 5A, B, C, F), WT or mutant HSP42 coding sequences (excluding stop codon) were cloned in pYM25 upstream and in frame with GFP using Gibson assembly. HSP42(WT or mutant)-GFP-hphNT2 was amplified with flanking DNA for integration designed to introduce HSP42-GFP downstream of HSP42 promoter at its native genomic locus in the hsp42Δ strain. For C-terminal labelling of Acc1 with mCherry at its native genomic locus (Figure 5E, F) mCherry-natNT2 was amplified from pBS35 + natNT2 plasmid with adequate flanking sequence for integration. Primer used for strains construction are listed in table S2. At each step, diploid cells were sporulated, and haploid spores were dissected on selection media and further sequencing and microscopic analysis confirmed integration. Culture from confirmed spores gave rise to homozygous diploid cells as these are homothallic spores. Competent cells were prepared and transformations performed using standard protocols (Amberg, Burke, and Strathern 2005). Yeast were grown in YPD medium containing 1% yeast extract (Bioshop), 2% peptone (Bioshop), and 2% glucose (Bioshop) with the appropriate antibiotic selection.

### Sporulation and germination

Sporulation was conducted on sporulation medium plates containing 1% potassium acetate, 0.1% yeast extract, 0.01% glucose, and 2% agar and spores were further purified on Percoll gradient (Sigma) as previously described (Plante and Landry 2020a). Germination was induced by transferring spores to YPD. To monitor germination, fresh spores were diluted in YPD at an OD_600_ = 1 and optical density was measured periodically in an Infinite M Nano plate reader (Tecan) set at 30°C.

### Heat shock resistance assay

Resistance measurements in spores during germination (Figure 1B) were conducted as described previously (Plante and Landry 2020a). Briefly, freshly purified wild spores were induced in germination in YPD medium, and at the indicated time following induction cells were sampled. Half of the cells were diluted in YPD medium, and the other half was treated at 55°C for 10 minutes in a thermocycler (Eppendorf Mastercycler ProS) before being transferred to YPD. Growth curves of both treated and untreated cells were recorded in an Infinite M Nano plate reader (Tecan) set at 30°C without shaking. Area under the curve (AUC) was determined using the Growthcurver package in R (Sprouffske and Wagner 2016). Heat resistance value was defined as the ratio of AUC of treated growth curve to AUC of untreated growth curve both obtained over the time required for untreated spore ODs to reach stationary phase. For resistance measurement of vegetative cells (Figure 5A), cells were grown overnight in YPD and diluted in YPD at OD_600_ of 0.1 and grown at 30°C until they reached an OD_600_ of 0.4 -0.5. Equal amounts of cell were diluted in fresh YPD medium, or incubated at 50°C for 10 minutes in a thermocycler prior to dilution. Growth curves of treated and control cells were recorded in a plate reader set at 30°C.

### Phase contrast and fluorescence cell imaging

All microscopic imaging experiments were performed using eight-well glass-bottom chamber slides (Sarstedt) coated with 0.05 mg/ml concanavalin A (Millipore Sigma). For phase contrast observation of germination (Figure 1C), freshly prepared spores were induced in germination by transferring them in a chamber filled with YPD medium. Cell imaging was performed on an Apotome Observer Z1 microscope (Zeiss) equipped with LD PlnN 40x/0.6 objective (Zeiss) at the indicated time after induction in a single field.

For fluorescence observation during germination, freshly prepared spores were diluted in YPD medium and incubated at 30 °C. At the indicated time after exposure to germination conditions, spores were washed in water, and transferred in a chamber filled with SC medium containing 0.174% Yeast nitrogen base (BioShop), 2% glucose and 0.5% ammonium sulfate (BioShop). For fluorescence observation on vegetative cells (Figure 5B), cells were grown in YPD at 30 °C until they reached an OD_600_ of 0.4 -0.5. Cells were left untreated (Control) or subjected to a heat shock at 50°C for 10 minutes in a thermocycler. They were then washed in water and transferred in a chamber filled with SD medium. Fluorescence imaging was performed on an Apotome microscope equipped with a Plan-Apochromate 100x/1.4 oil objective (Zeiss). Image acquisition was performed using an AxioCam MRm camera (Zeiss). Images were analysed using the ImageJ software (Schneider, Rasband, and Eliceiri 2012).

### Fluorescence heterogeneity score

To measure cellular heterogeneity score, fluorescent images were opened in ImageJ and cell peripheries were drawn by hand to analyse fluorescence signals of single cells. Heterogeneity scores are the fluorescence signal covariant coefficients, obtained by dividing the standard variation by the mean signal.

### Transmission Electron microscopy

Freshly prepared spores were induced in germination in YDP at 30°C. At the indicated time after the induction of germination, cells were harvested, washed in water and suspended in fixative solution, containing 2.5% glutaraldehyde, 1.5% paraformaldehyde, 0.5mM CaCl_2_ in 0.1M caco buffer pH 7.2. Vegetatively growing cells in YPD (OD600 = 0.5-0.6) were harvested, washed in water and suspended in fixative solution. Cells were fixed for 24 hours at room temperature. Following steps were conducted by the microscopy platform of IBIS (Université Laval, Québec, Canada).

Cells were dehydrated with ethanol solution (30-100%), then embedded in epoxy resin (Epon). 150 nm–thick sections of resin-embedded cells were prepared using an ultramicrotome (Ultracut UCT; Leica), and stained with 1% (wt/vol) uranyl acetate in 70% (wt/vol) methanol for 5 min and 0.4% lead citrate for 3 min. Samples were imaged on a JEM 1230 Transmission Electron Microscope (JOEL). Images were analysed and processed using ImageJ software.

### Molecular probes

The complete sequence of mammalian orthoreovirus 3 strain T3 non structural protein µNS (GeneBank MK246417.1) was kindly shared by Pr. Martin Bisaillon from Université de Sherbrooke. We cloned by gibson assembly the whole coding sequence, minus stop codon, into pYM25 (Janke et al. 2004) to generate a fusion with yeGFP at its C-terminus. The promoter of *SOD1* (nucleotides -851 to -1 relative to ATG) was cloned by Gibson assembly upstream the µNS CDS in pYM25. Expression of SOD1 was shown in spores and during germination (Plante et al. 2017), and expression of the molecular probe with this promoter happened at a high level in spores and during germination which suited our experiments with this cell type. *SOD1* promoter - µNS - GFP in addition to HPH markers on pYM25 were amplified as a whole with the appropriate flanking sequences for genomic integration at the *URA3* locus. From all tested loci for integration (*MET15*, *LEU2*, *HIS3*), *URA3* allows high and uniform expression of the probes across the population, while having the least effect on sporulation and germination efficiency.

Plasmid p426MET25 containing sfpHluorin gene was purchased from Addgene (ID 115697). We swapped the yeGFP gene in pYM25 plasmid for sfpHluorin, and cloned the *SOD1* promoter upstream the sfpHluorin CDS by Gibson assembly. The *SOD1* promoter - sfpHluorin in addition to HPH marker on pYM25 were amplified as a whole with the appropriate flanking sequences for genomic integration at the *URA3* locus. Yeast with either genomic integration were selected for hygromycin resistance.

### Particle tracking and microrheology

Cells expressing *µNS*-GFP were transferred to a 8-well glass-bottom chamber slides (Sarstedt) coated with concanavalin A 0.05 mg/mL (Millipore Sigma) and filled with 500µl of complete SC medium. Image acquisition was performed using a Perkin Elmer UltraVIEW confocal spinning disk unit attached to a Nikon Eclipse TE2000-U inverted microscope equipped with a Plan Apochromat DIC H 100×/1.4 oil objective (Nikon), and a Hamamatsu Orca Flash 4.0 LT + camera. Imaging was done at 30 °C in an environmental chamber. The software NIS-Elements (Nikon) was used for image capture. For each field, one brightfield and a series of fluorescence (GFP) images were taken. Cells were excited with a 488 nm laser and emission was filtered with a 530/630 nm filter. GFP time lapse images were acquired continuously at a rate of two frames/sec for one min. Images were processed using image J. Particle tracking was performed using the python package Trackpy ((Crocker and Grier 1996), http://soft-matter.github.io/trackpy/v0.5.0/). Particles were identified in microscopic images using the “locate” function. Minimal mass threshold was set at 200 to exclude spurious fluorescence signals. Trajectories were assembled from the multiple frames using the “link” function. The “imsd” and “emsd’ function was used to compute mean squared displacement of individual particles and ensemble MSD respectively. Microns per pixel was set as 10/75 and frames per second = 2.

### Intracellular pH measurements

Exponentially growing wild type cells expressing sfpHluorin (OD = 0.3-0.4) in YPD medium were used for calibration curve determination as previously described (Triandafillou and Drummond 2020). Cells were washed twice in water and suspended in calibration buffer containing 50 mM NaCl, 50 mM KCl, 50 mM MES, 50 mM HEPES, 100 mM ammonium acetate, 10 mM 2- deoxyglucose and 10 µM nigericin; pH was adjusted with HCl or KOH from 5.0 to 9.0. After 30 minutes incubation at room temperature, fluorescence (533 nm) of sfpHluorin following excitation at 405 and 488 nm was acquired using a Guava EasyCyte HT cytometer (EMD Millipore). The calibration curve was generated by taking the median ratio of fluorescence after excitation at 405 nm to excitation at 488 nm (405/488 ratio) at various pH. Ratios were corrected for background by subtracting the autofluorescence of unlabeled cells (WT). Points were fitted to a sigmoid (Figure S1D). Viscosity is among many parameters that affect the response of pHluorin to its environment (Fricker et al. 2001). To confirm that pHluorin sensitivity and response in the highly dense and rigid ascospores cytosol is comparable to that in yeast cells, we performed a calibration curve in spores. Spores expressing pHluorin were incubated in a calibration buffer adjusted to various pH for 1 hour at room temperature before fluorescence was analyzed by flow cytometry as for vegetative yeast. Fluorescence ratios showed that pHluorin in spores have a similar sensitivity and response compared to that in yeast cells (Figure S1D). pH measurement was performed on vegetatively growthing cells (OD=0.3-0.4) expressing sfpHluorin in YPD and freshly prepared spores expressing sfpHluorin at the indicated time-points after exposure to rich medium. Cells were washed twice in water then suspended in a measurement buffer containing 50 mM NaCl, 50 mM KCl, 50 mM MES, 50 mM HEPES, 100 mM ammonium acetate. After 30 minutes of incubation at room temperature, the median 405/488 ratio was measured by cytometry. pH values were obtained from the sigmoid function of the calibration curve.

### Protein extraction and sedimentation

Freshly purified wild spores at the indicated time following germination induction in YPD medium, and vegetatively growing cells in YPD (OD = 0.5-0.6) were harvested. Cell were resuspended in 4 ml Protein buffer containing 120 mM KCl, 2 mM EDTA, 20 mM HEPES-KOH, pH 7.4, 1:500 Protease inhibitor (MiliporeSigma), 0.5 mM DTT and 1mM PMSF, and snap frozen as 20 µL beads, then placed in a 10 ml milling pod (Retsch) cooled in liquid nitrogen along with a 10 mm milling bead. 20 milling cycles of 2 minutes each each were performed on a Mixer Mill MM 400 (Retsch) at 30 Hz, with cooling in liquid nitrogen between each cycle. Cell extracts were thawed on ice, and clarified by centrifugation at 16,000 g for 10 minutes. Supernatant was retrieved and protein concentration was measured by BCA protein assay (Novagen, (Smith et al. 1985). Protein concentrations were adjusted in all the samples to 800 µg/ml. Equal volume (2 ml *i.e* 1600 µg) of cell extracts were loaded in ultracentrifuge tubes (Beckman). Samples were ultracentrifuged at 100,000 g for 30 minutes at 4°C in a Optima XPN-100 ultracentrifuge (Beckman). Supernatants were kept aside as “Supernatant” fraction. Pellets were washed twice with protein buffer, then resuspended in protein buffer + 1% SDS which correspond to “Pellet” fraction. 1% of total supernatant (i.e. 20 µl) and pellet fraction (i.e. 10 µl) for each cell extract was loaded on a 10% SDS-polyacrylamide gel in a loading buffer containing 0.06M Tris pH 6.8, 0.07M SDS, 10% glycerol, 5% mercaptoethanol and 0.01% bromophenol blue. Migration was conducted at 90 V until dye front reached 1 cm into the gel. Proteins were stained with Coomassie G-250 dye, and lanes were cut out of the gel and stored in 1.5 ml microtubes before they are further processed.

### Western Blot

For detection of native proteins in fractionated cell extracts (Figure S3), 1% of supernatant and pellet fraction for each time-point in germination were loaded on a 12% SDS-polyacrylamide gel. Migration was conducted at 120 V until dye front reach the bottom of separating gel. Proteins were then transferred to a nitrocellulose membrane (Li-Cor), and blocked for 2 hours at room temperature in a blocking Buffer (Li-Cor). The following antibodies were used for detection of Bcy1, Homocitrate synthase and actin, respectively: anti-Bcy1 (yN-19, Santa Cruz Biotechnology, SC-6764), Homocitrate Synthase (31F5, Santa Cruz Biotechnology, SC-57832), and anti-actin (Clone C4, EMD Milipore, MAB1501R). After washing in phosphate buffered saline (PBS) 1x containing 1% Tween 20, membranes were incubated with the appropriate fluorophore-conjugated antibodies (Li-Cor). Blots were then imaged on an Odyssey Imager (Li-Cor), and images were analysed on Image Studio software (Li-Cor, v1.1).

### Mass spectrometry

In gel protein digestion was performed as previously described (Shevchenko et al. 1996). Gel lanes of each sample were cut into smaller pieces, destained with 40% ethanol in 30 mM ammonium bicarbonate then reduced with 10 mM DTT at 37°C for 30 min then alkylated with 55 mM iodoacetamide at 37°C for 30 min. The gel pieces were digested at 37°C initially with 0.5 µg of trypsin (Promega) per sample for 6 hours then additionally with 0.3 μg of trypsin overnight. The resulting peptides were extracted from gel pieces using sequential shaking in 40% acetonitrile then 100% acetonitrile, vacuum centrifuged (Vacufuge, Eppendorf) to evaporate the organic solvents and cleaned through C18 STop-And-Go-Extraction tips (StageTips, PMID 12498253), eluted in 40% acetonitrile, 0.1% formic acid, and vacuum centrifuged until complete dryness.

LC-MSMS analysis (Kerr et al. 2020). The concentration of the final reconstituted sample was measured at *A*_205_ using a NanoDrop One (Thermo Fisher) to inject 250 ng into Bruker Impact II Qtof coupled to easy nLC 1200 (Kerr et al. 2020). The injection was randomized to minimize loading order bias. A single analytical column set up using IonOpticks’ Aurora UHPLC column (1.6 μm C18 and 25 cm long) was used to create 90 minutes of separation from 5% to 35% buffer B for each sample.

### Data search

Resulting data were searched on MaxQuant version 1.6.17.0 (Cox et al. 2009) against sequences from verified and uncharacterized ORFs from the R64-3-1 release of the S288C genome proteome database (yeastgenome.org) and common contaminant sequences provided by the software (246 sequences) adding the following variable modifications: oxidation on methionines, acetylation on protein N-termini, acetylation on lysines, methylations on arginine, and phosphorylation on serines, threonines, and tyrosines. Fixed carbamidomethylation was set on cysteines. Default match between runs was enabled and default peptide and fragment mass tolerances (10 and 40 ppm) were set. Data were filtered to have 1% false discovery rates at peptide and protein levels. The mass spectrometry proteomics data have been deposited to the ProteomeXchange Consortium via the PRIDE (Perez-Riverol et al. 2022) partner repository with the dataset identifier PXD035403.

### Proteomic analysis

For the analysis of protein solubility (Pindex measurements) we considered the intensity-based absolute quantification (iBAQ, (Schwanhäusser et al. 2011)) of proteins with sequence coverage of ≥10% with at least 2 peptides. 895 proteins, for which total abundance (Supernatant + pellet) was > 0 in each replicate at every time-points of germination, were included in the analysis. Pindex of a given protein was measured as the ratio of its abundance in the pellet to its total abundance (Supernatant + Pellet).

Since Pindex values across the triplicates were highly correlated (Figure S2A), we considered the mean Pindex of each triplicate. Pindex values of the 895 proteins considered at each time-point in germination are listed in Table S3. Clustering of Pindex trajectories was performed in python using the Hierarchical clustering method in the Scipy package (scipy.cluster.hierarchy). Hierarchical linkage was conducted with the linkage function using the “complete” method. The clusters were then defined using the fcluster function using the “distance” criterion for discrimination.

### Protein properties

Molecular weight and isoelectric point of the 895 considered proteins were retrieved on web-based YeastMine application (https://yeastmine.yeastgenome.org/). Total iBAQ (Supernatant + Pellet) for each of the 895 proteins in the analysis was average amongst the five time-points to obtain the mean abundance. To measure, in the considered proteins, the amino acid composition predicted to form prion-like domain, we used the Prion-like amino acid composition (PLAAC) web-based application (http://plaac.wi.mit.edu/details, (Lancaster et al. 2014)). From this application, we considered the normalized score (NLLR) of each protein for our analysis. To predict the propensity of each of the proteins to condensate, we used the python application PSAP ((van Mierlo et al. 2021), https://github.com/Guido497/phase-separation). This classifier scores each residue, and we used the median score of each protein for further analysis. To predict the consensus disorder of each protein, we used the python application Metapredict ((Emenecker, Griffith, and Holehouse 2021), https://github.com/idptools/metapredict) which is a neural network trained for single residue scoring. For further analysis, we used the median metapredict score for each protein. Principal component analysis (PCA) was performed using the Scikit-learn package in python. Protein properties data were first scaled using the StandardScaler function, then PCA was performed with PCA function.

### Trehalose quantification

Trehalose content was assayed following a method by (Chen and Futcher 2017). Equal number (10^8^) of spores at the indicated time after exposure to rich medium or exponentially growing yeast (vegetative) were harvested and cleaned with water.

Carbohydrates were then extracted through an alkali treatment. Cells were incubated in 0.25M Na_2_CO_3_ at 95°C for 3 hours. After incubation, pH was adjusted to 5.8 by addition of 2.5 volume of freshly prepared 0.2M NaOAc. The extract was then separated in half. Add 1% of porcine trehalase (Sigma, T8778) and incubate overnight at 37°C to allow hydrolysis of trehalose in glucose. Glucose concentration was assayed in the extracts using a commercial kit (Sigma, GAGO20). Trehalose concentration in extracts was measured as the glucose concentration after trehalase treatment, minus glucose concentration without treatment.

## Supplemental information

**Figure S1.**
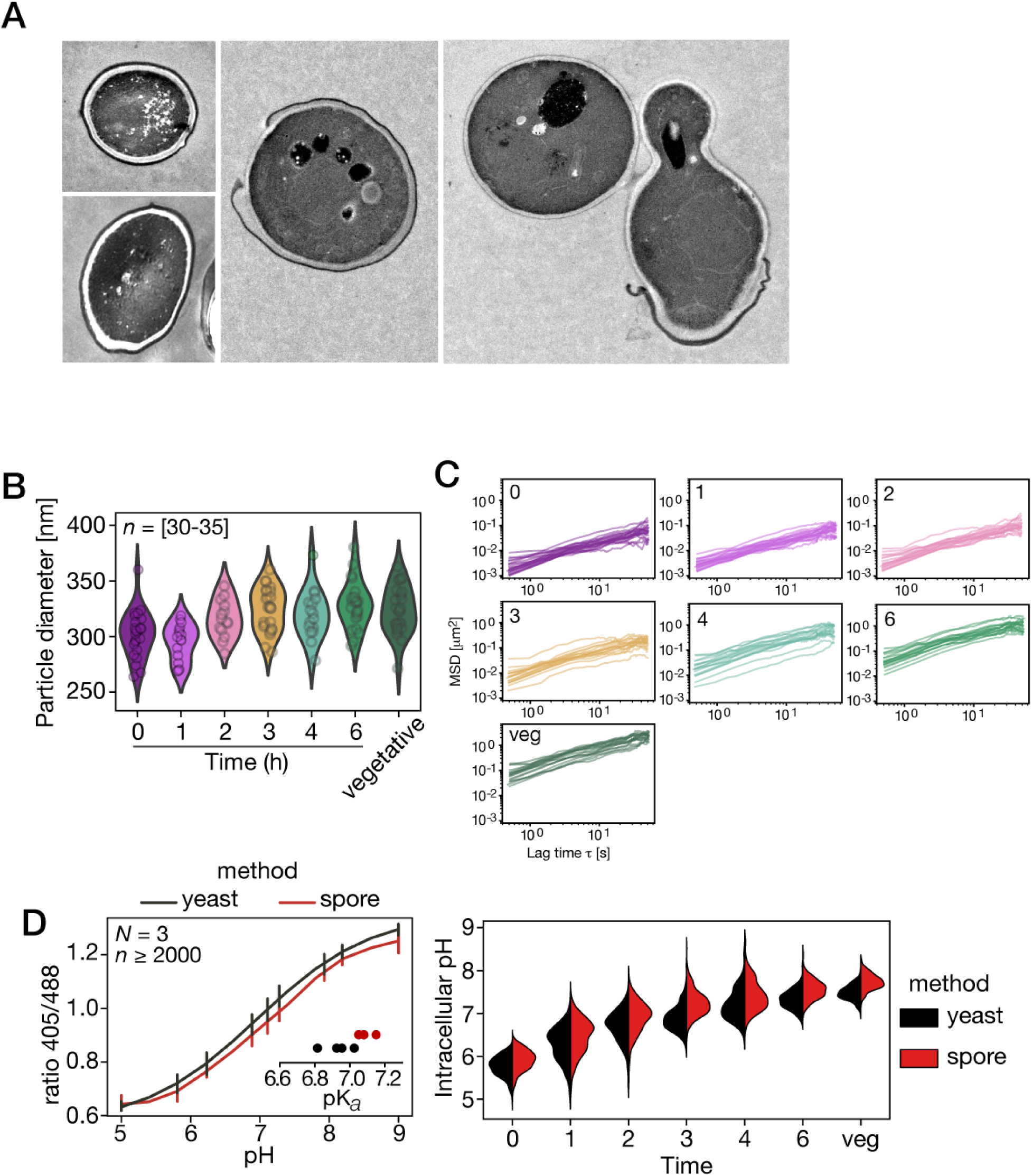
µNS-GFP particle size and Mean square displacement (MSD) during germination; calibration curve of pHluorin emission ratio, related to figure 1. A) Transmission electron microscopy images of spores at the indicated time after exposure to rich medium. Scale bar represents 1 µm. B) Size of individual particles tracked at each time point during germination and in vegetatively growing cells. At least 30 particles were tracked at each time point. C) Mean square displacement of individual particles tracked at the indicated time after germination induction. D) Left, intracellular pH calibration curves determined using vegetative yeast (black) or spores (red). Although the curve using spores is slightly more basic, measurement of pH in germination (right) using either curve shows that spores are acidic and that intracellular pH increases steadily during germination. Logistic function fitted to the data from vegetative cells was used for Figure 1G. Error bars represent standard deviation of three replicates.

**Figure S2.**
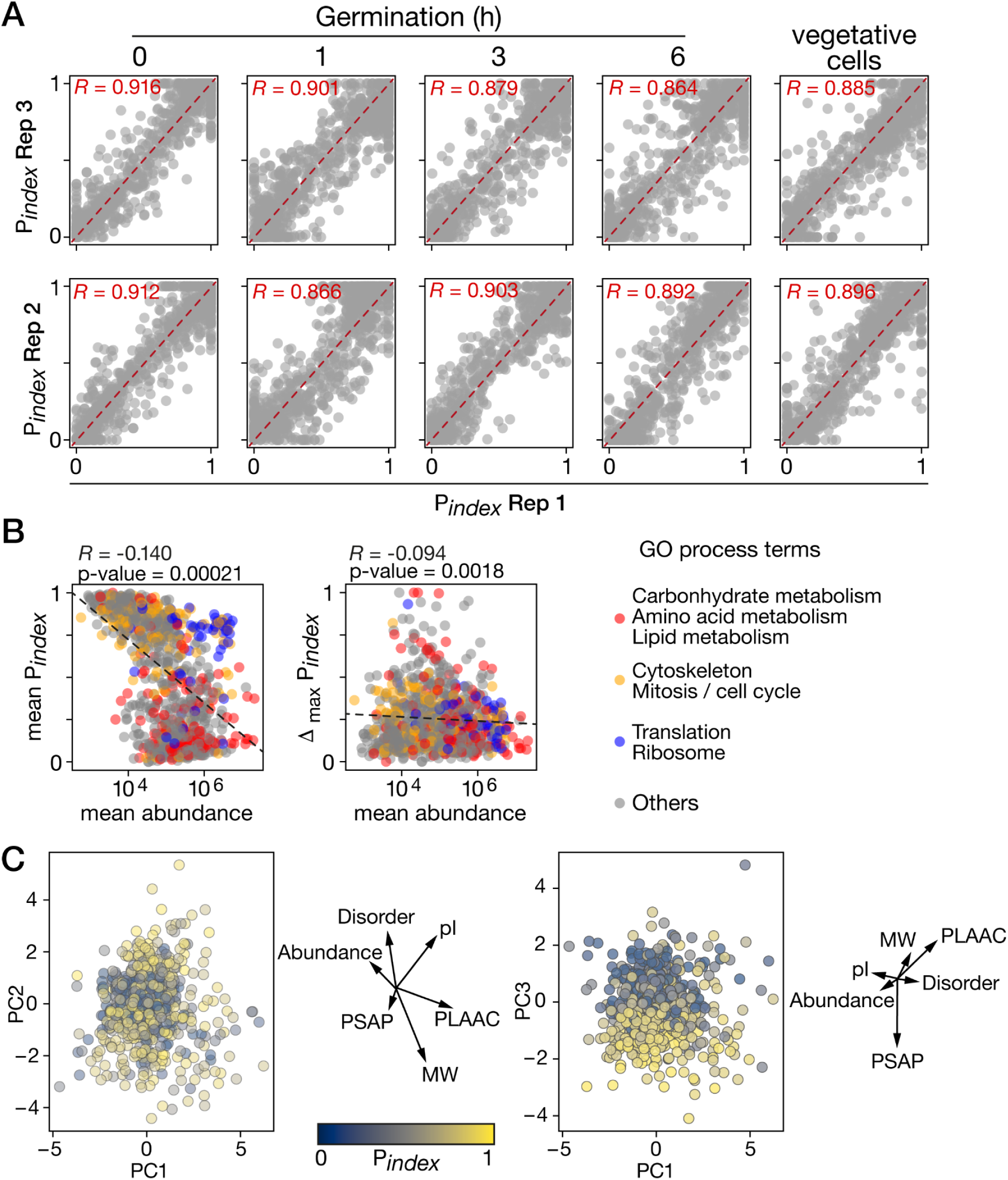
Pindex correlation between replicates, influence of protein abundance on solubility measurements, and contribution of protein properties on P_index_ distribution. Related to Figure 2. A) P_index_ values are plotted against other replicates. Pearson’s correlation coefficients are indicated on each graph. For all correlations, p-value < 0.0001. B) Mean of the absolute abundance estimated from mass spectrometry data during germination is plotted against mean Pindex values (left), or the maximal Pindex variation (right, Δ_max_P_index_) of each protein. Points are colored depending on the GO function term. Pearson’s correlation coefficient with the log10-transformed abundance values are shown with the corresponding p-values. C) PCA analysis of protein properties. Protein distribution across PC1 vs PC2 (left) and PC1 vs PC3 (right). Dots are colors according to the mean P_index_ value. Beside the graph is the vector representation indicating the strength and direction of the contribution of each variable to the distribution; sequence-based estimation of molecular weight (MW) and isoelectric point (pI); mean abundance measured from our proteomic data; Prion-like amino acid composition (PLAAC) prediction score; analysis and prediction score of phase separation (PSAP); sequence-based prediction of disorder (Metapredict)

**Figure S3.**
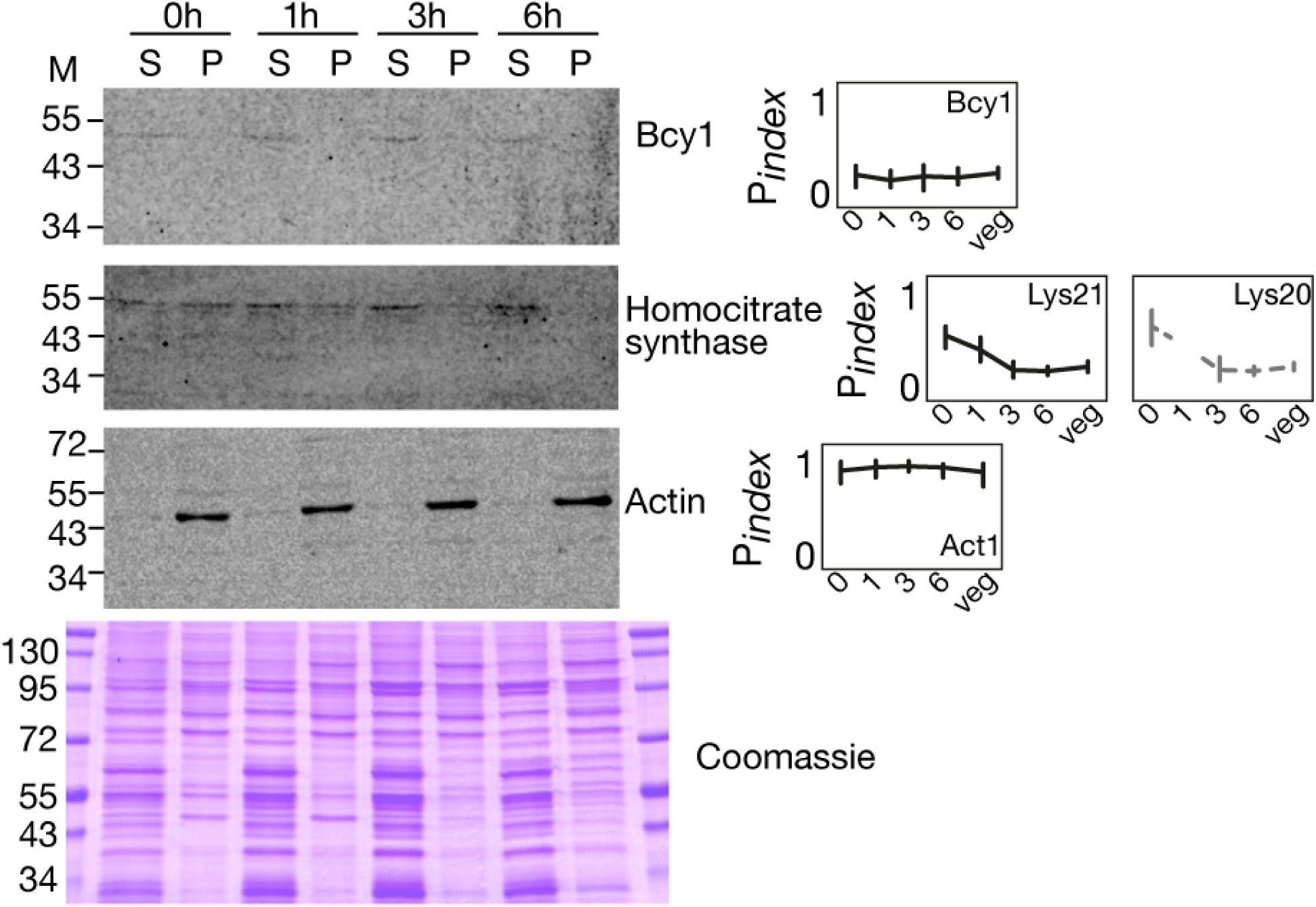
Protein sedimentation reveals different protein solubility, related to Figure 2. Left, Same fractionned protein extract (S, supernatant fraction; P, pellet fraction) used in MS measurements were analysed by SDS-PAGE with antibodies that were available for yeast endogenous proteins. Shown are anti-Bcy1, anti-Homocitrate synthase and anti-actin western blots, and an identically loaded gel stained by Coomassie. Proteins molecular weight in the ladder (NEB# P7706) are indicated in KDa. Right, P*index* trajectories of the proteins analyzed by western blot. Homocitrate synthase isozyme Lys20 was poorly detected at 1 hour time-point. Error bars represent standard deviation of three replicates.

**Figure S4.**
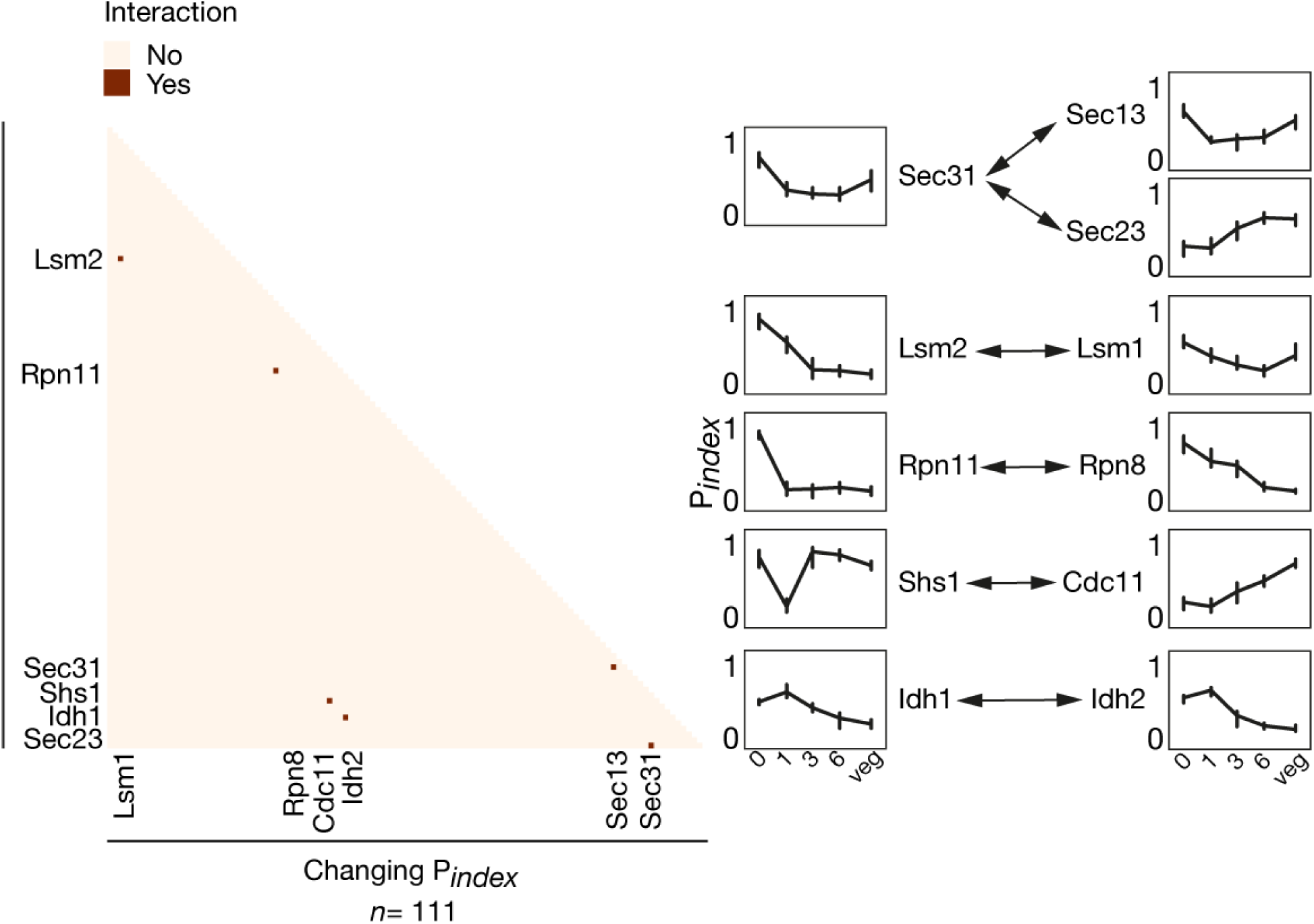
Known protein-protein interaction among proteins of the changing Pindex group. Left, interaction between the 111 Changing Pindex proteins were searched through the known physical interaction database (BioGRID v4.4.216). Pairs of interacting proteins are marked in red in the heat map with the identity of the partners on the side. Right, the 6 pairs of interacting proteins are indicated with arrows, and the Pindex trajectories for each protein is shown. Error bars represent standard deviation of three replicates.

**Figure S5.**
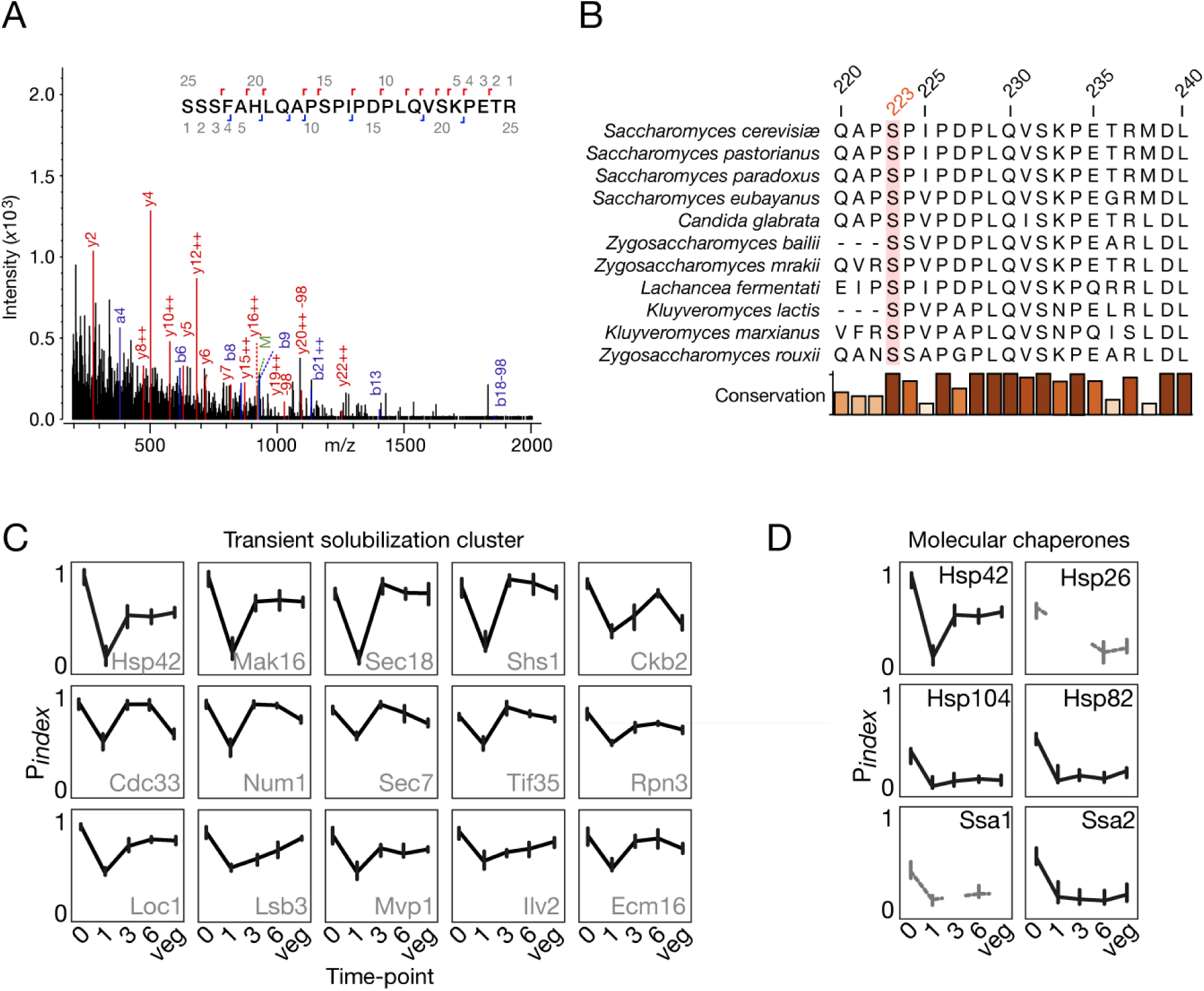
Phosphorylation of Hsp42 on S223, Transient solubilization cluster and molecular chaperones, related to Figure 4. A) MS spectra example of phosphorylated S223 peptide on Hsp42. B) Multiple Sequence Alignment of Hsp42 orthologs. Numbers on top refer to residue position in the *S. cerevisiae* protein. S223 is underlined in orange. Relative conservation is shown with the bars at the bottom. Only a small portion of the sequences are shown. C) Individual P*index* trajectories for each 15 proteins in the transient solubilization cluster. Error bars represent standard deviation of three replicates. D) P*index* trajectories of molecular chaperones detected in our experiments. Hsp26 and Ssa1 have only partial data since they are not well detected. However, these data reveal that Hsp42 have a unique sedimentation profile among molecular chaperones. Error bars represent standard deviation of three replicates.

**Figure S6.**
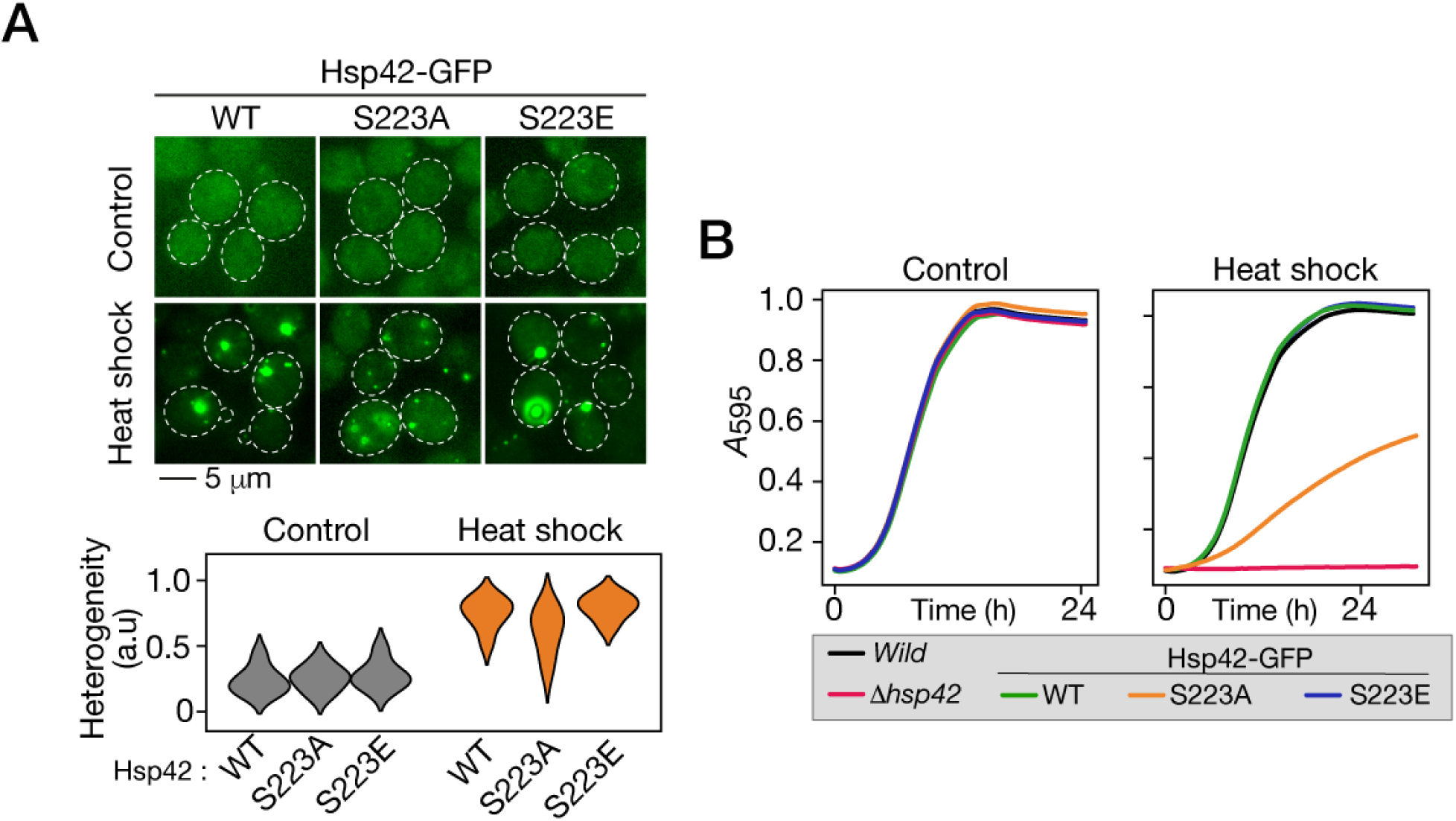
Stress activation of Hsp42 through its phosphorylation on S223, related to Figure 5. A) Fluorescence microscopic images of WT or mutant Hsp42-GFP expressing cells, in control conditions (top) or after a heat shock (bottom). Dotted lines represent cell contour. Scale bar represents 5 µm. Bottom, cellular Hsp42-GFP heterogeneity measure. The S223A Hsp42 mutant shows smaller and fainter aggregates in cells, and lower heterogeneity score compared to WT and phosphomimetic mutant. Heat shock at 50°C for 10 minutes. B) Growth curves of vegetative cells of the indicated strains after a heat shock (right) or a mock treatment at control temperature (left). Shown are the mean values of three replicates. The S223A Hsp42 mutant shows intermediate heat-shock resistance between the WT (and S223E mutant) and *HSP42* deleted cells. This confirms that the phosphorylation of Hsp42 at this site is important for its function.

**Table S1:** Key resource table

| REAGENT or RESOURCE | SOURCE | IDENTIFIER |
| --- | --- | --- |
| **Chemicals, peptides, and recombinant proteins** |  |  |
| cOmplete, EDTA-free Protease Inhibitor Cocktail | MiliporeSigma | cat#11836153001 |
| Percoll | MiliporeSigma | cat#P1644 |
| Nigericin | MiliporeSigma | cat#481990 |
| 2-Deoxyglucose | Bioshop | cat#DXG498 |
| Concanavalin A | MiliporeSigma | cat#C2010 |
| **Critical commercial assays** |  |  |
| BCA Protein Assay Kit | Novagen | Cat#71285 |
| **Deposited data** |  |  |
| Raw and analyzed mass spectrometry data | Data are available via ProteomeXchange | PXD035403 |
| **Experimental models: *Saccharomyces cerevisiae* strains** |  |  |
| LL13_054 wild diploid strain MATa/$\alpha$ | (Leducq et al. 2016) | LL13_054 |
| ura3::PSOD1-μNS-GFP hphNT1 (background: LL13_054) | This Paper | SPY020 |
| ura3::PSOD1-sfpHluorin hphNT1 | This Paper | SPY031 |
| (background:<br>LL13_054) |  |  |
| ACC1-GFP::hphNT<br>1<br>(background:<br>LL13_054) | This Paper | SPY037 |
| URA7-GFP::hphNT<br>1<br>(background:<br>LL13_054) | This Paper | SPY039 |
| HSP42-GFP::hphNT<br>1<br>background:<br>LL13_054) | This Paper | SPY040 |
| GLK1-GFP::hphNT1<br>(background:<br>LL13_054) | This Paper | SPY044 |
| hsp42Δ::KanMX4<br>(background:<br>LL13_054) | This Paper | SPY056 |
| hsp42Δ::HSP42-GF<br>P-hphNT1<br>(background:<br>LL13_054) | This Paper | SPY078 |
| hsp42Δ::HSP42(S2<br>23A)-GFP-hphNT1<br>(background:<br>LL13_054) | This Paper | SPY080 |
| hsp42Δ::HSP42(S2<br>23D)-GFP-hphNT1<br>(background:<br>LL13_054) | This Paper | SPY081 |
| ACC1-mCherry::nat<br>NT2<br>(background:<br>LL13_054) | This Paper | SPY089 |
| hsp42Δ::KanMX4<br>ACC1-mCherry::nat<br>NT2<br>(background:<br>LL13_054) | This Paper | SPY093 |
| hsp42Δ::HSP42-GF<br>P-hphNT1<br>ACC1-mCherry::nat<br>NT2<br>(background:<br>LL13_054) | This Paper | SPY101 |
| hsp42Δ::HSP42(S2<br>23A)-GFP-hphNT1<br>ACC1-mCherry::nat<br>NT2<br>(background:<br>LL13_054) | This Paper | SPY102 |
| hsp42Δ::HSP42(S2<br>23D)-GFP-hphNT1<br>ACC1-mCherry::nat<br>NT2<br>(background:<br>LL13_054) | This Paper | SPY103 |
| **Oligonucleotides** |  |  |
| Primers used in this<br>study are listed in<br>table S1 | This study | N/A |
| **Recombinant DNA** |  |  |
| Plasmid: pYM25 | PCR tool box | Janke et al., Yeast, 2004 |
| plasmid:<br>pYM25-PSOD1-μN<br>S-yeGFP | This study | N/A |
| plasmid:<br>pYM25-PSOD1-sfp<br>Hluorin | This study | N/A |
| plasmid: pUG6 | Euroscarf | P30114 |
| plasmid: pNATCRE | (Steensma and Ter Linde 2001) | pNATCRE |
| plasmid: pBS35 (mCherry) + natNT2 | Addgene | Cat#83797 |
| plasmid: pYM25-HSP42-GFP | This study | N/A |
| plasmid: pYM25-HSP42(S22 3A)-GFP | This study | N/A |
| plasmid: pYM25-HSP42(S22 3D)-GFP | This study | N/A |
| **Software and algorithms** |  |  |
| Rstudio | Rstudio | RRID: SCR_000432 (https://www.rstudio.com/) |
| Python (v 3.7.4) | Python | https://www.python.org/ |
| GrowthCurver (v 0.3.1) | (Sprouffske and Wagner 2016) | https://github.com/sprouffske/growthcurver |
| TrackPy (v 0.5.0) | (Crocker and Grier 1996) | http://soft-matter.github.io/trackpy/v0.5.0/ |
| Matplotlib (v 3.5.1) | (Hunter 2007) | https://matplotlib.org/ |
| Seaborn | (Waskom 2021) | https://seaborn.pydata.org/index.html |
| Prion-Like Amino Acid Composition (PLAAC) | (Lancaster et al. 2014) | http://plaac.wi.mit.edu |
| scipy.cluster.hierarchy (v1.8.1) | Scipy | https://docs.scipy.org/doc/scipy/reference/cluster.hierarchy.html |
| Metapredict (v2.0) | (Emenecker, Griffith, and Holehouse 2021) | https://github.com/idptools/metapredict |
| Phase Separation Analysis and Prediction (PSAP) | (van Mierlo et al. 2021) | https://github.com/Guido497/phase-separation. |
| Scikit-learn (sklearn) v1.1.1 | (Pedregosa et al. 2011) | https://scikit-learn.org/ |

**Table S2.**
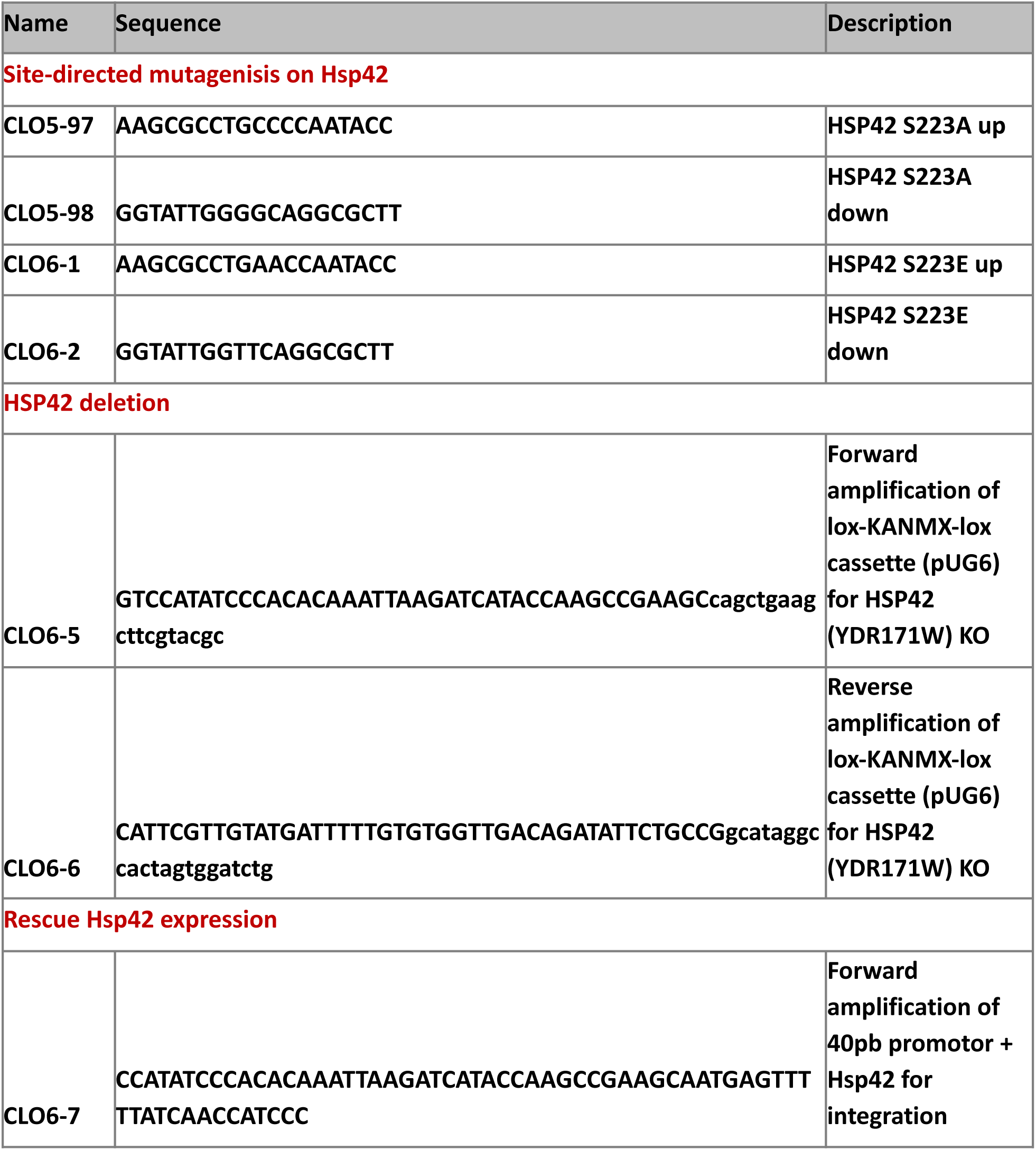
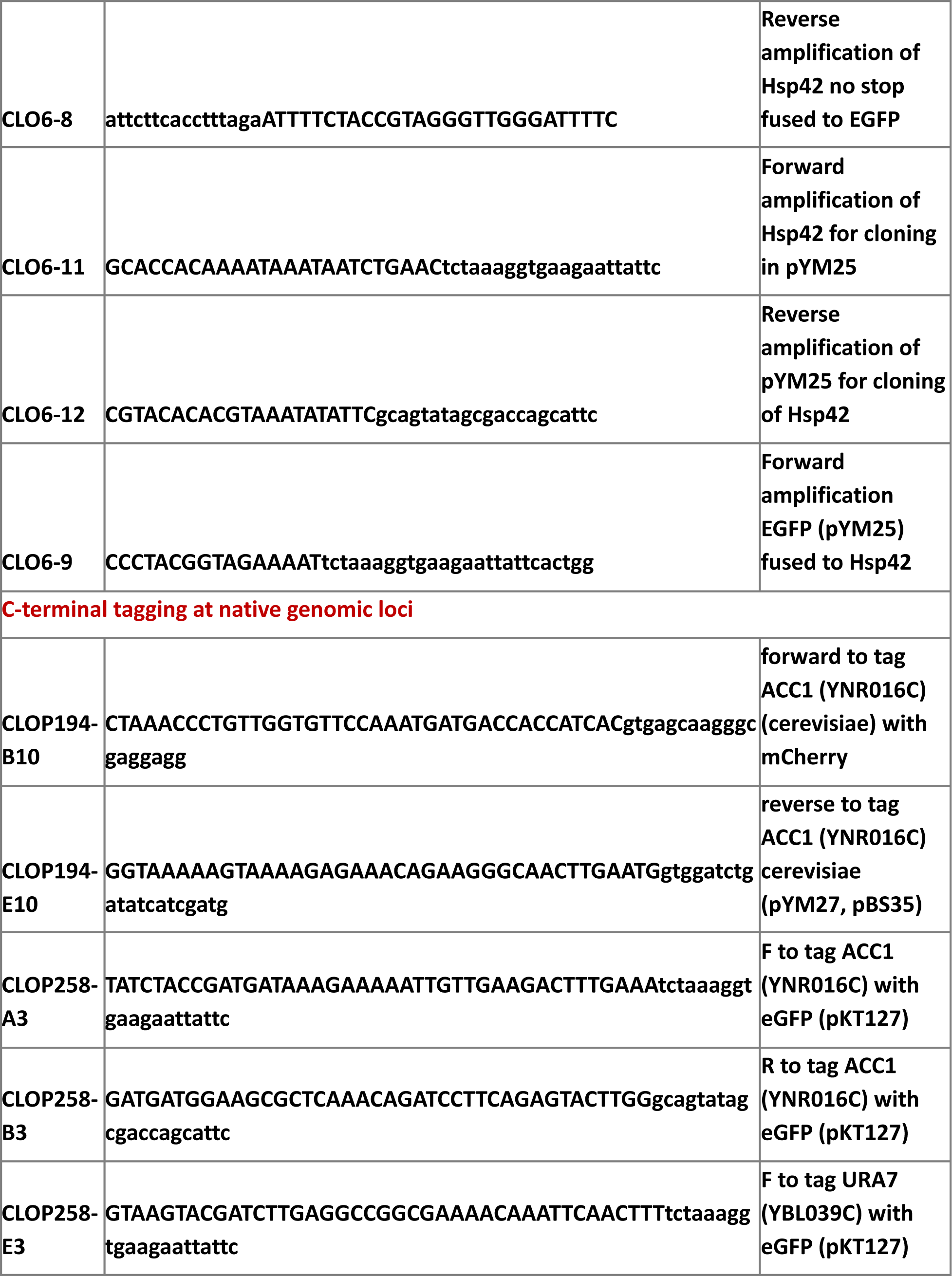
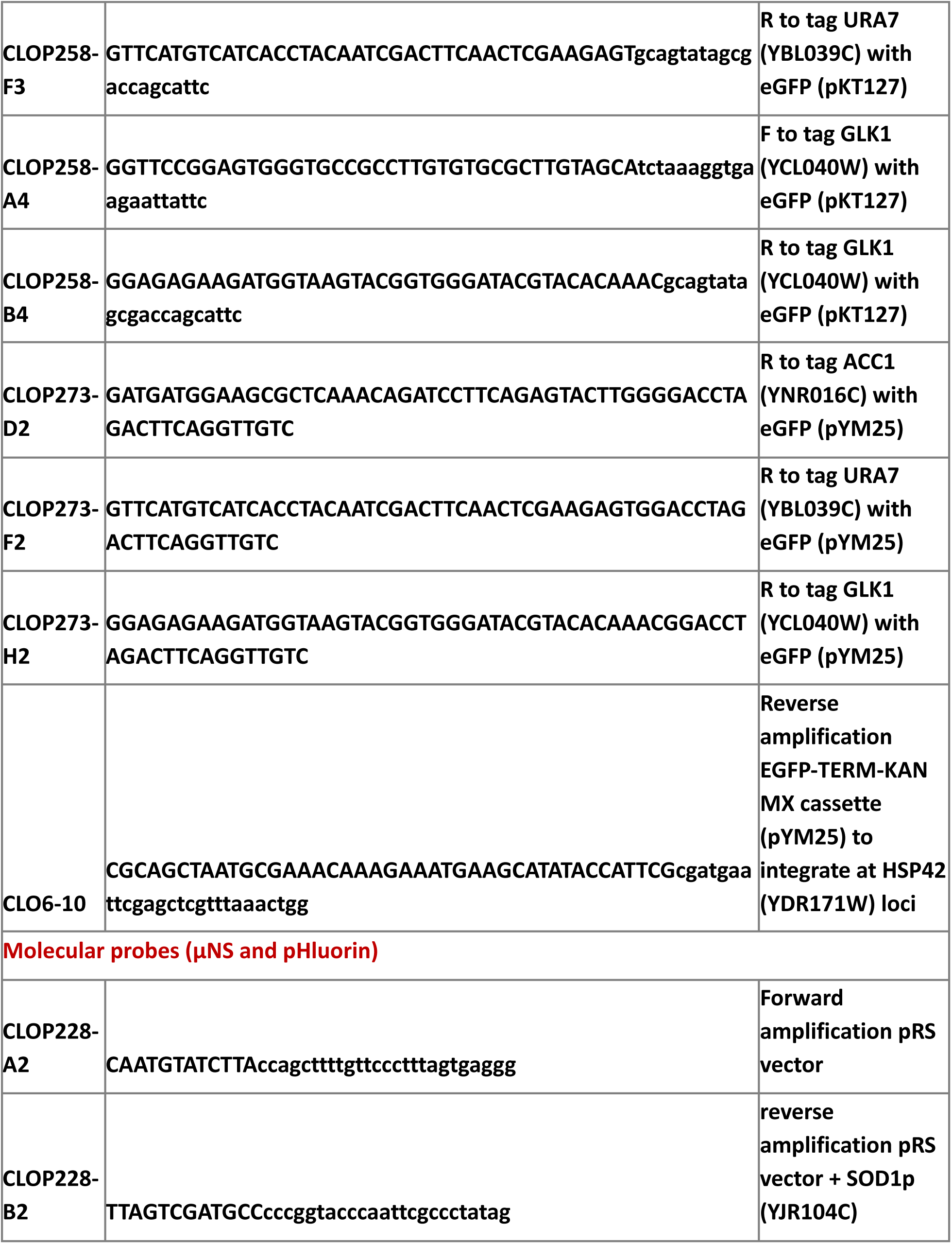
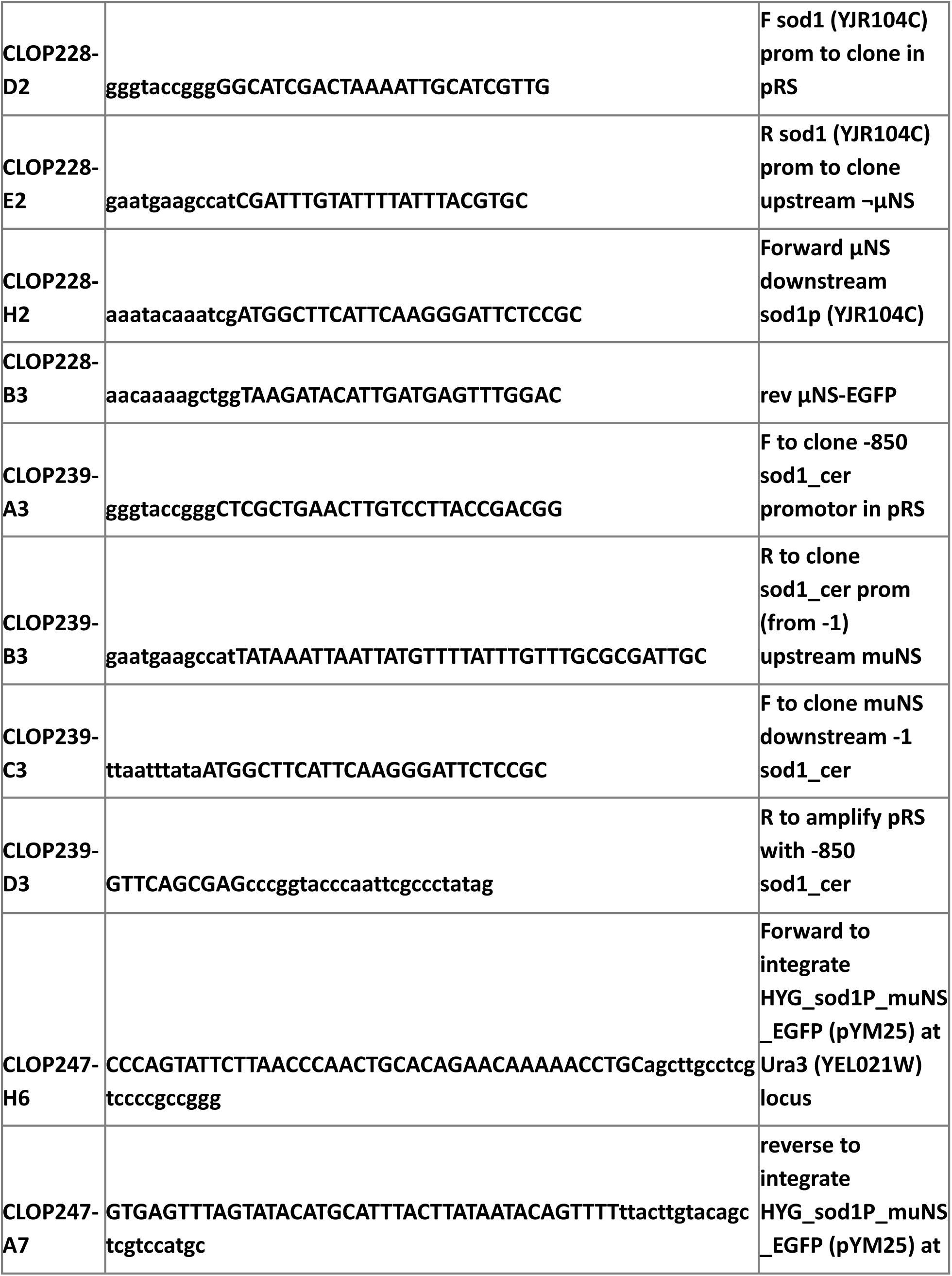
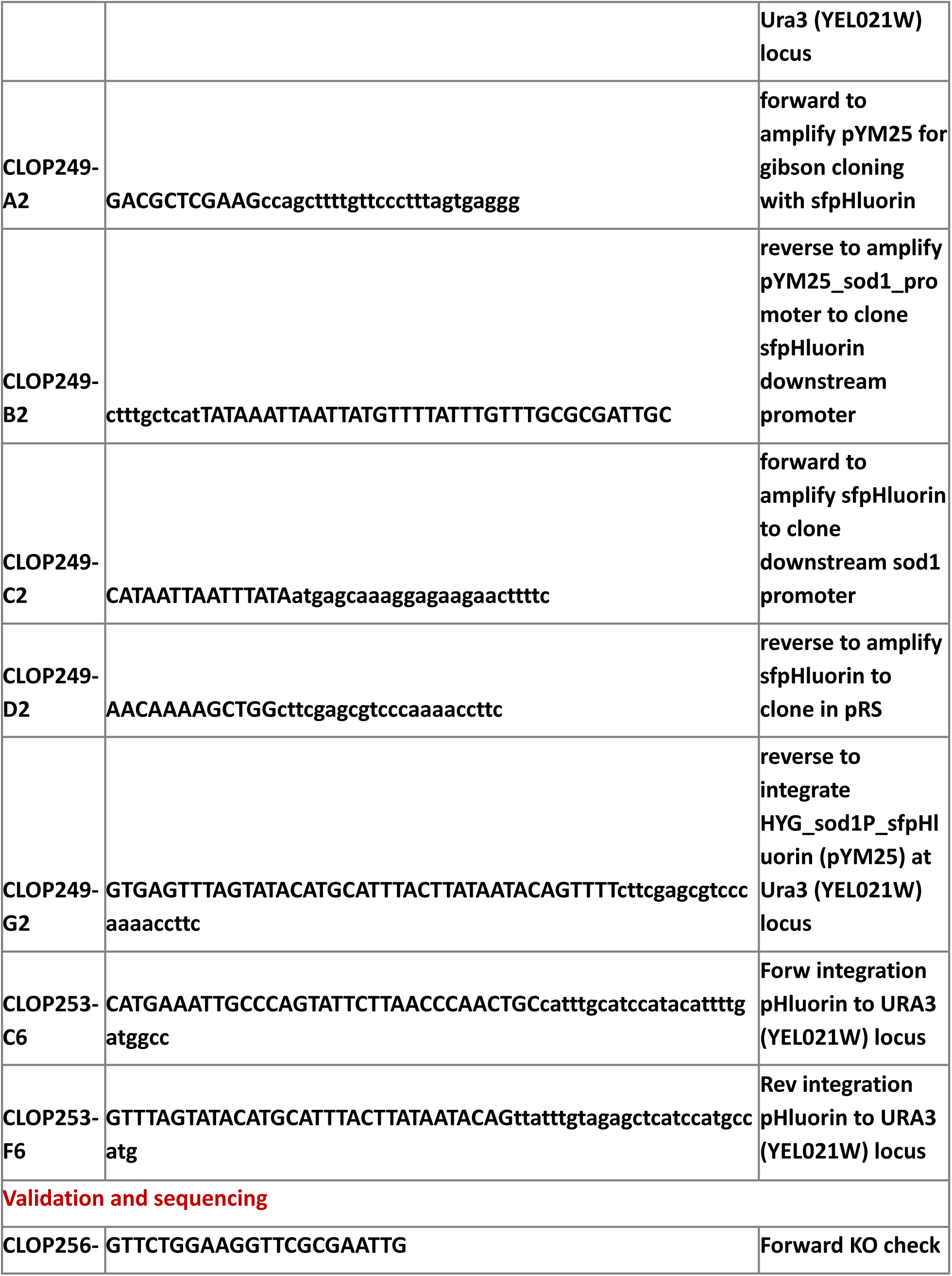
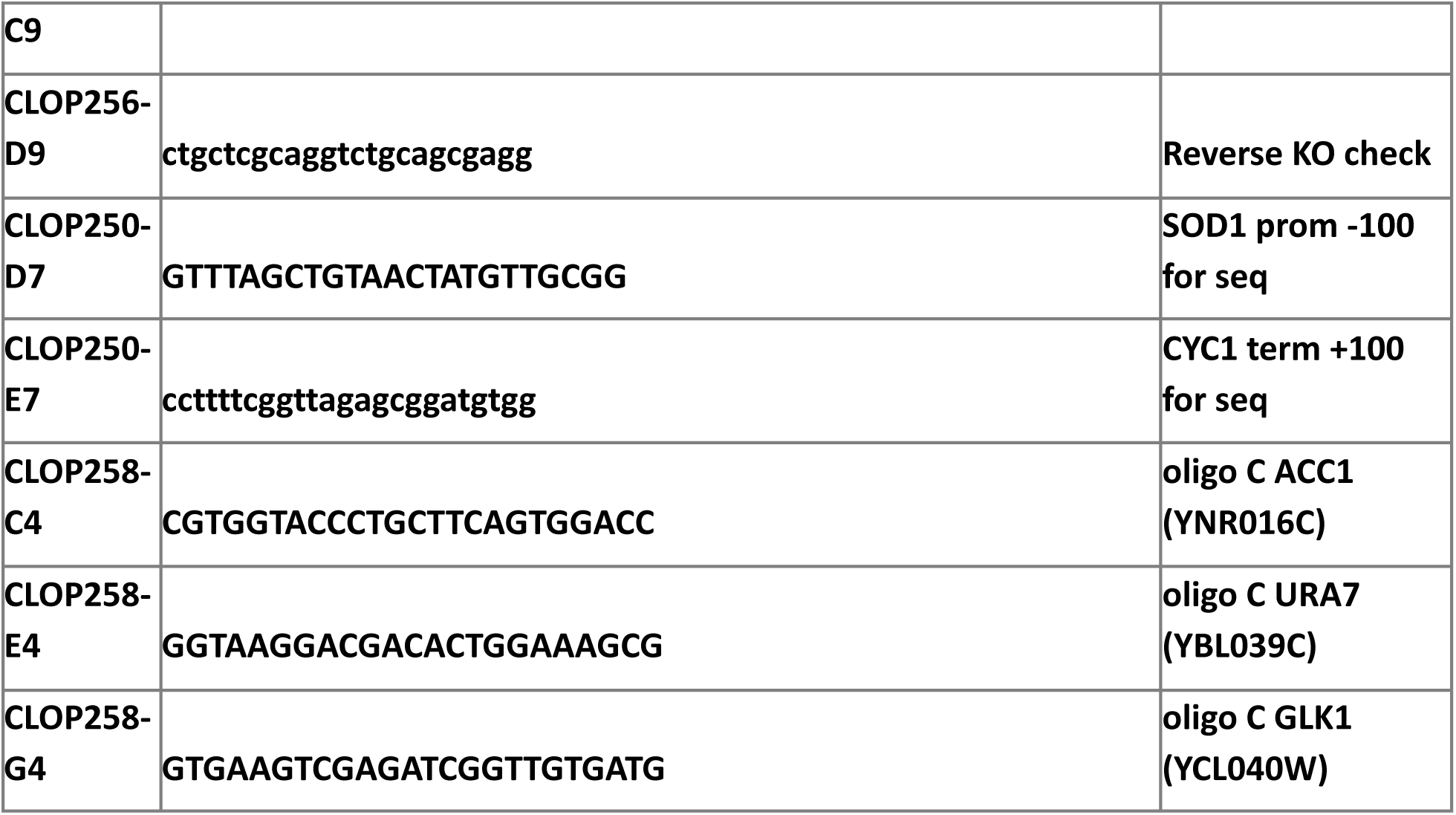
Primers used in this study

**Table S3.** Pindex values at indicated time-point during germination and in vegetatively growing cells.

