## Supplementary material for "Breaking dormancy in spores of budding yeast transforms its cytoplasm and the solubility of its proteome": Table S1

Key resource table S1

| REAGENT or RESOURCE | SOURCE | IDENTIFIER |
| --- | --- | --- |
| Chemicals, peptides, and recombinant proteins | | |
| cOmplete, EDTA-free Protease Inhibitor Cocktail | MiliporeSigma | cat#11836153001 |
| Percoll | MiliporeSigma | cat#P1644 |
| Nigericin | MiliporeSigma | cat#481990 |
| 2-Deoxyglucose | Bioshop | cat#DXG498 |
| Concanavalin A | MiliporeSigma | cat#C2010 |
| Critical commercial assays | | |
| BCA Protein Assay Kit | Novagen | Cat#71285 |
| Deposited data | | |
| Raw and analyzed mass spectrometry data | Data are available via ProteomeXchange | PXD035403 |
| Experimental models: *Saccharomyces cerevisiæ* strains | | |
| LL13_054 wild diploid strain MATa/𝞪 | (Leducq et al., 2016) | LL13_054 |
| ura3::P*SOD1*-µNS-GFP hphNT1  (background: LL13_054) | This Paper | SPY020 |
| ura3::P*SOD1*-sfpHluorin hphNT1  (background: LL13_054) | This Paper | SPY031 |
| ACC1-GFP::hphNT1  (background: LL13_054) | This Paper | SPY037 |
| URA7-GFP::hphNT1  (background: LL13_054) | This Paper | SPY039 |
| HSP42-GFP::hphNT1  background: LL13_054) | This Paper | SPY040 |
| GLK1-GFP::hphNT1  (background: LL13_054) | This Paper | SPY044 |
| hsp42∆::KanMX4  (background: LL13_054) | This Paper | SPY056 |
| hsp42∆::HSP42-GFP-hphNT1  (background: LL13_054) | This Paper | SPY078 |
| hsp42∆::HSP42(S223A)-GFP-hphNT1 (background: LL13_054) | This Paper | SPY080 |
| hsp42∆::HSP42(S223D)-GFP-hphNT1  (background: LL13_054) | This Paper | SPY081 |
| ACC1-mCherry::natNT2  (background: LL13_054) | This Paper | SPY089 |
| hsp42∆::KanMX4 ACC1-mCherry::natNT2  (background: LL13_054) | This Paper | SPY093 |
| hsp42∆::HSP42-GFP-hphNT1 ACC1-mCherry::natNT2  (background: LL13_054) | This Paper | SPY101 |
| hsp42∆::HSP42(S223A)-GFP-hphNT1 ACC1-mCherry::natNT2  (background: LL13_054) | This Paper | SPY102 |
| hsp42∆::HSP42(S223D)-GFP-hphNT1 ACC1-mCherry::natNT2  (background: LL13_054) | This Paper | SPY103 |
| Oligonucleotides | | |
| Primers used in this study are listed in table S1 | This study | N/A |
| Recombinant DNA | | |
| Plasmid: pYM25 | PCR tool box | Janke et al., Yeast, 2004 |
| plasmid: pYM25-PSOD1-µNS-yeGFP | This study | N/A |
| plasmid: pYM25-PSOD1-sfpHluorin | This study | N/A |
| plasmid: pUG6 | Euroscarf | P30114 |
| plasmid: pNATCRE | (Steensma and Ter Linde, 2001) | pNATCRE |
| plasmid: pBS35 (mCherry) + natNT2 | Addgene | Cat#83797 |
| plasmid: pYM25-HSP42-GFP | This study | N/A |
| plasmid: pYM25-HSP42(S223A)-GFP | This study | N/A |
| plasmid: pYM25-HSP42(S223D)-GFP | This study | N/A |
| Software and algorithms | | |
| Rstudio | Rstudio | RRID: SCR_000432 (https://www.rstudio. com/) |
| Python (v 3.7.4) | Python | https://www.python.org/ |
| GrowthCurver (v 0.3.1) | (Sprouffske and Wagner, 2016) | https://github.com/sprouffske/growthcurver |
| TrackPy (v 0.5.0) | (Crocker and Grier, 1996) | http://soft-matter.github.io/trackpy/v0.5.0/ |
| Matplotlib (v 3.5.1) | (Hunter, 2007) | https://matplotlib.org/ |
| Seaborn | (Waskom, 2021) | https://seaborn.pydata.org/index.html |
| Prion-Like Amino Acid Composition (PLAAC) | (Lancaster et al., 2014) | http://plaac.wi.mit.edu |
| scipy.cluster.hierarchy (v1.8.1) | Scipy | https://docs.scipy.org/doc/scipy/reference/cluster.hierarchy.html |
| Metapredict (v2.0) | (Emenecker et al., 2021) | https://github.com/idptools/metapredict |
| Phase Separation Analysis and Prediction (PSAP) | (van Mierlo et al., 2021) | https://github.com/Guido497/phase-separation). |
| Scikit-learn (sklearn) v1.1.1 | (Pedregosa et al., 2011) | https://scikit-learn.org/ |
